# TAF4b transcription networks regulating early oocyte differentiation

**DOI:** 10.1101/2021.07.18.452838

**Authors:** Megan A. Gura, Sona Relovska, Kimberly M. Abt, Kimberly A. Seymour, Tong Wu, Haskan Kaya, James M. A. Turner, Thomas G. Fazzio, Richard N. Freiman

## Abstract

Establishment of a healthy ovarian reserve is contingent upon numerous regulatory pathways during embryogenesis. Previously, mice lacking TBP-associated factor 4b (*Taf4b*) were shown to exhibit a diminished ovarian reserve. However, potential oocyte-intrinsic functions of TAF4b have not been examined. Here we use a combination of gene expression profiling and chromatin mapping to characterize the TAF4b gene regulatory network in mouse oocytes. We find that *Taf4b*-deficient oocytes display inappropriate expression of meiotic, chromatin, and X-linked genes, and unexpectedly we found a connection with Turner Syndrome pathways. Using Cleavage Under Targets and Release Using Nuclease (CUT&RUN), we observed TAF4b enrichment at genes involved in meiosis and DNA repair, some of which are differentially expressed in *Taf4b*-deficient oocytes. Interestingly, TAF4b target genes were enriched for Sp/KLF family motifs rather than TATA-box, suggesting an alternate mode of promoter interaction. Together, our data connects several gene regulatory nodes that contribute to the ovarian reserve.

## INTRODUCTION

The ability to produce healthy gametes is critical for the continuation of all sexually reproducing organisms, including humans. The process of mammalian gametogenesis begins during early fetal life with the specification and migration of primordial germ cells (PGCs) to the genital ridge. At the genital ridge, PGCs in close concert with sex-specific somatic support cells begin the process of differentiation into eggs and sperm. Thus, to understand the healthy functioning of adult gametes, we must examine multiple stages of development including those that arise in early fetal life. An added layer of complexity is that female XX and male XY germ cells traverse this differentiation process in a highly sex-specific manner (Feng et al. 2014). While some adult male germ cells become self-renewing spermatogonial stem cells (SSCs) during their development, the adult female mammalian germline is a non-renewable and finite resource termed the ovarian reserve that is steadily depleted after birth. The postnatal ovarian reserve is composed of a stockpile of primordial follicles (PFs) that contain individual primary oocytes arrested in prophase I of meiosis I, surrounded by a single layer of flattened somatic granulosa cells (Gura and Freiman 2018). Menopause results from the timely depletion of the ovarian reserve and the mean age for menopause is 50±4 years. At least 1% of the female population worldwide experiences a fertility deficit termed primary ovarian insufficiency (POI) where menopause-like symptoms occur prematurely by 40 years of age (Chandra et al. 2013). Thus, genetic and environmental factors that perturb the establishment of the ovarian reserve *in utero* will have negative consequences on adult reproductive and general health outcomes and need to be understood in greater detail.

We previously identified an essential function of TBP-Associated Factor 4b (TAF4b) in the establishment of the ovarian reserve in the embryonic mouse ovary (Grive et al. 2016, 2014). TAF4b is a germ cell-enriched subunit of the transcription factor TFIID complex, which is required for RNA Polymerase II recruitment to promoters in gonadal tissues (Gura et al. 2020). TFIID is a multi-protein complex that contains TATA-box binding protein (TBP) and 13-14 TBP-associated factors (TAFs) and is traditionally considered part of the cell’s basal transcription machinery (Antonova et al. 2019). Male mice that have a targeted mutation, which disrupts the endogenous *Taf4b* gene and prevents TAF4b protein from integrating into the larger TFIID complex (called *Taf4b*-deficiency), are subfertile. *Taf4b*-deficient female mice are infertile and also exhibit hallmarks of POI including elevated follicle stimulating hormone (FSH) levels and a diminished ovarian reserve (DOR) (Falender et al. 2005; Gura et al. 2020; Lovasco et al. 2015, 2010a). We recently demonstrated that *Taf4b* mRNA and protein expression are nearly exclusive to the germ cells of the mouse embryonic ovary from embryonic day 9.5 (E9.5) to E18.5 and that *Taf4b*-deficient ovaries display delayed germ cell cyst breakdown, increased meiotic asynapsis, and excessive perinatal germ cell death (Grive et al. 2016, 2014; Gura et al. 2020).Therefore, we hypothesize that TAF4b, as part of TFIID, regulates oogenesis and meiotic gene programs. To what degree the transcriptomic pathways in *Taf4b*-deficiency and POI overlap and contribute to their similarities has yet to be explored.

Both human and mouse genetic studies have begun to reveal the molecular mechanisms underlying POI and its related pathologies. The most striking example is Turner Syndrome (TS) where karyotypically single X-chromosome female individuals undergo early and severe DOR and experience short stature, primary amenorrhea, estrogen insufficiency, and cardiovascular malformations (Gravholt et al. 2019). Recent work in mouse models of TS indicates that loss of correct dosage of the single X chromosomes in XO versus XX oocytes leads to pronounced meiotic progression defects and excessive oocyte attrition as the ovarian reserve is being established (Sangrithi et al. 2017). In contrast to the high penetrance of TS, 20% of women with a premutation CGG repeat allele in the *FMR1* gene, also located on the X chromosome, experience a related fragile X-associated POI (FXPOI) (Fink et al. 2018). Similar to *Taf4b*, other targeted mouse mutations have resulted in POI-related phenotypes including those in *Nobox* and *Figla*, two transcription factors that regulate oocyte development, however the relevance of specific mutations in their human orthologs and POI in women remains to be explored (Rossetti et al. 2017). More importantly, a better understanding of how these genes promote healthy establishment of the ovarian reserve and the deregulated molecular events that lead to its premature demise is needed.

To better understand the normal function of TAF4b during establishment of the ovarian reserve, we integrated published bioinformatic data with experimental *Taf4b* genomic assays to uncover unexpected links of *Taf4b* with TS and *Fmr1*. We show that in homozygous mutant *Taf4b* E16.5 oocytes, almost 1000 genes are deregulated as measured by RNA-sequencing (RNA-seq). Surprisingly, the X chromosome was enriched for these deregulated genes and *Taf4b*-deficient oocytes display reduced X:autosome (X:A) gene expression ratios. There is a striking overlap of genes deregulated in *Taf4b*-deficient oocytes and XO mouse oocytes, and XO oocytes express significantly reduced levels of *Taf4b* at E15.5 and E18.5, further illuminating a potential molecular link between these disparate genetic contributors to POI. Further, we show that *Taf4b*-deficiency and TS both result in deregulation of genes involved in chromatin organization, modification, and DNA repair. Finally, CUT&RUN of TAF4b E16.5 XX and XY germ cells identifies direct TAF4b targets that for the first time confirm its promoter-proximal recognition properties, linking TAF4b binding to the critical transcriptional regulation required for the proper establishment of the ovarian reserve.

## RESULTS

### Taf4b expression peaks at E16.5 in female embryonic germ cells

To observe the dynamics of *Taf4b* mRNA expression at a single cell resolution, we analyzed a single-cell RNA-seq (scRNA-seq) dataset of Oct4-GFP-positive oocytes from E12.5, E14.5, and E16.5 mouse ovaries (Zhao et al. 2020). We selected for *Dazl*-positive, high quality (nFeature_RNA > 1000, nFeature_RNA < 5000, nCount_RNA > 2500, nCount < 30000, percent mitochondrial genes < 5%) oocytes and performed pseudotime analysis using Monocle3 (**Fig 1A-B**). We found that *Figla* expression generally increased over time and pseudotime, with some of the highest Figla-expressing cells appearing in E16.5 cells at the end of the pseudotime profile and *Stra8* expression, which is a master regulator of meiotic initiation, declined over time as expected. We then compared the expression profiles of *Taf4a* and *Taf4b*. Most cells across the time course had low *Taf4a* expression throughout. *Taf4b* mRNA expression began to rise at E14.5 and appeared highest in the E16.5 oocytes that were earliest in pseudotime (**Fig 1C**), consistent with previous observations (Gura et al. 2020).

**Figure 1.**
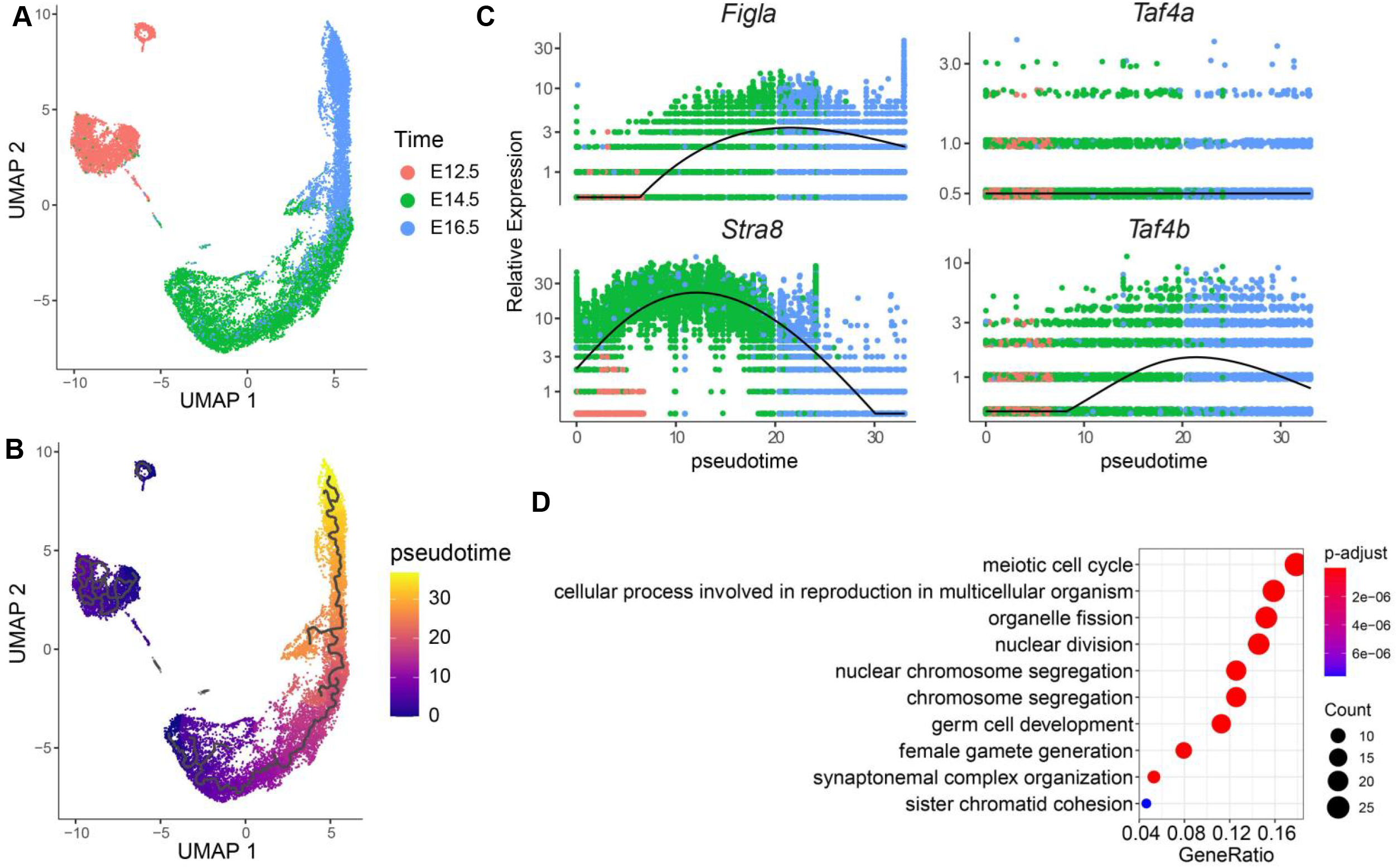
Analysis of scRNA-seq dataset in E12.5 to E16.5 germ cells. (A) Uniform Manifold Approximation and Projection (UMAP) of oocytes colored by embryonic time point. (B) UMAP of oocytes colored by pseudotime analysis. (C) Expression of *Figla*, *Stra8*, *Taf4b*, and *Taf4a* plotted in terms of pseudotime and colored based on embryonic time point. (D) Dotplot of GO results for genes significantly higher in *Taf4b*-expressing oocytes than *Taf4b*-non-expressing oocytes.

To identify which other genes were highly expressed in *Taf4b*-expressing oocytes, we performed differential gene expression analysis using cells separated into *Taf4b*-expressing (*Taf4b* log_2_ expression > 0) and *Taf4b*-off (*Taf4b* log_2_ expression = 0) populations and performed differential gene expression analysis (**Table S1**). We found 155 genes significantly (p-value < 0.05) higher in *Taf4b*-expressing cells. We performed gene ontology (GO) analysis of these genes and found that the top categories included “meiotic cell cycle” and “synaptonemal complex organization” (**Fig 1D, Table S1**). Taken together, these data suggest that *Taf4b* expression is highest in the E16.5 mouse oocyte and that *Taf4b* is co-expressed with important meiotic genes. A similar analysis from a second scRNA-seq dataset of whole ovaries from earlier E11.5 to E14.5 time points supported these findings (Ge et al. 2021) (**Fig S1**).

### RNA-seq identifies Taf4b-affected genes in E16.5 XX germ cells

To understand the transcriptome-level changes in *Taf4b*-deficient embryonic oocytes, we performed RNA-seq at E16.5. We sorted Oct4-GFP-positive oocytes from five *Taf4b*-heterozygous (*Taf4b* +/-) and five *Taf4b*-deficient (*Taf4b* -/-) pairs of ovaries and subjected them to ultra-low input RNA-seq. The resulting principal component analysis (PCA) plot shows each of the *Taf4b*-deficient samples mostly grouping together, with the *Taf4b*-heterozygous samples dispersed throughout (**Fig 2A, Table S2**). This patterning of the data is largely due to the litter from which each sample originates, as we were unable to obtain sufficient numbers of our desired genotypes from a single mouse litter, but importantly the different genotypes separate when plotting litter dates individually (**Fig S2A**). We identified 964 differentially expressed genes (DEGs) between *Taf4b*-heterozygous and *Taf4b*-deficient oocytes, which were defined as protein-coding, average transcripts per million (TPM) expression > 1, and adjusted p-value < 0.05 (**Fig 2B, Table S2**). From this list of DEGs, 463 were increased in *Taf4b*-deficient oocytes and will be referred to as “Up in Def DEGs”. Some interesting DEGs in this gene set were *Fmr1* (the most common genetic cause of POI) and *Mki67* (a marker of cell proliferation) (**Fig S2B**). From the DEG list, 501 were decreased in *Taf4b*-deficient oocytes and will be referred to as “Down in Def DEGs”. As expected, *Taf4b* was a Down in Def DEG, as were other more well-known oogenesis genes *Sohlh1, Nobox*, and *Ddx4* (**Fig S2C**). Analysis of known protein-protein interactions (PPIs) using STRING revealed a significant enrichment of PPIs, with major nodes including *Ep300* (a histone acetyltransferase involved in chromatin remodeling) and *Plk1* (a serine/threonine-protein kinase involved in cell cycle regulation) (**Fig S3**). We also performed a similar RNA-seq experiment at E14.5, but found fewer DEGs suggesting that more substantial transcriptomic effects of *Taf4b*-deficiency take place around E16.5 (**Fig S4A-B, Table S3**).

**Figure 2.**
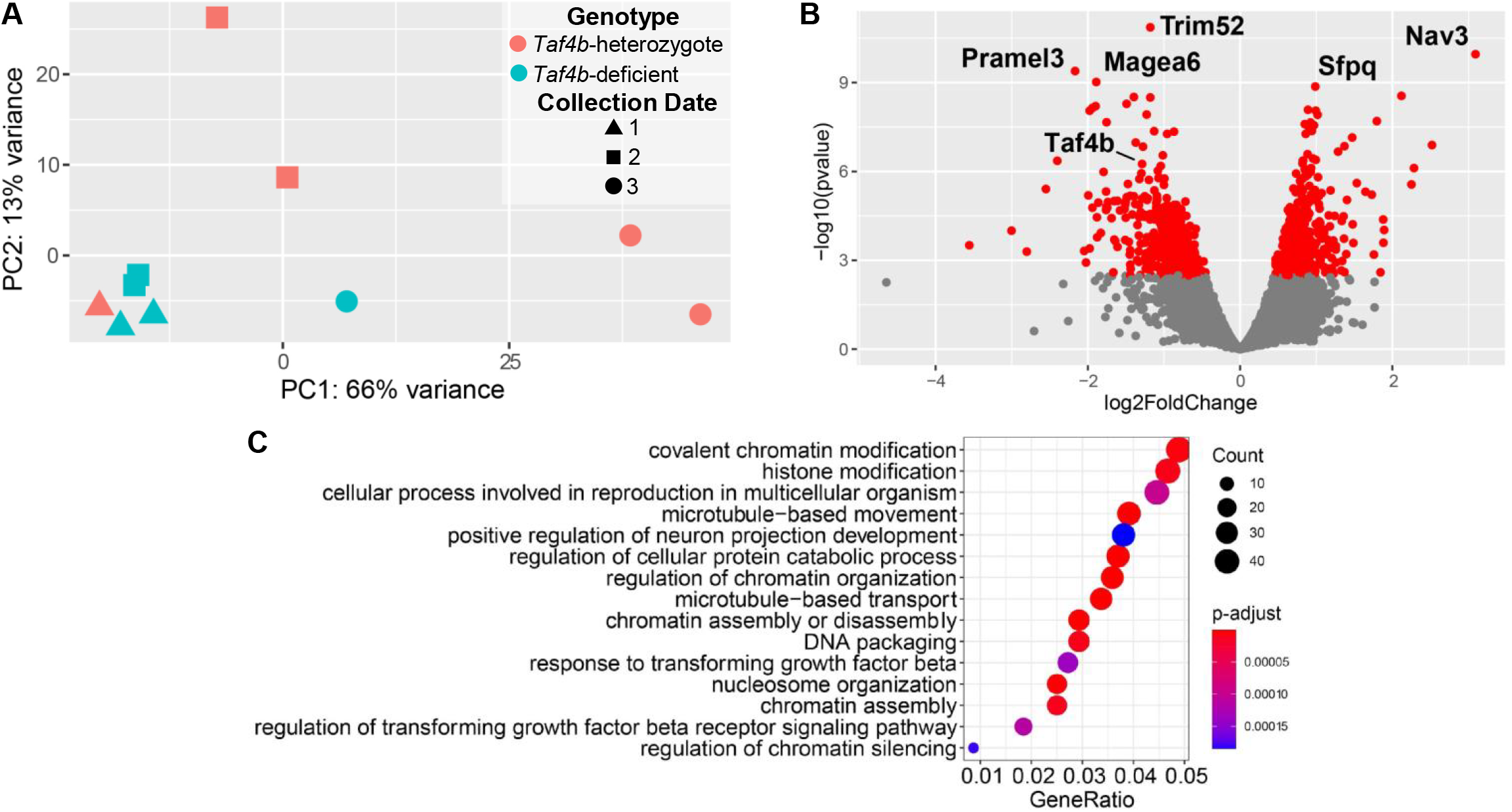
RNA-seq of E16.5 oocytes. (A) PCA plot of the E16.5 samples labeled based on **Taf4b** genotype and collection number. (B) Volcano plot of genes, the significant genes (protein-coding, p-adj < 0.05, avg TPM > 1) are labeled in red and the top 5 DEGs plus **Taf4b** are specified. (C) Dotplot of GO biological process analysis of DEGs.

We performed GO analysis of all the E16.5 DEGs, as well as separating the Up in Def and Down in Def DEGs (**Fig 1C; Fig S2D-E**). We found multiple chromatin organization and modification GO categories associated with Up in Def DEGs and reproduction- and microtubule-related categories associated with Down in Def DEGs (**Table S2**). Overall, these data suggest that *Taf4b* impacts the expression of many genes in the developing oocyte transcriptome, particularly those associated with chromatin structure and modification and reproduction. Moreover, the effects of *Taf4b* on the transcriptome take place after E15.5, correlated with the peak in *Taf4b* expression at E16 shown by scRNA-seq and bulk RNA-seq comparisons.

### X chromosome gene expression is significantly reduced in *Taf4b*-deficient oocytes

Our E16.5 RNA experiment led us to examine how *Taf4b*-deficiency affects expression of each mouse chromosome. Surprisingly, we observed that there were significantly (p < 0.05) more Down in Def DEGs on the X chromosome than expected and significantly (p < 0.05) fewer Up in Def DEGs on the X chromosome (**Fig 3A-B, Table S4-5**). Furthermore, the X chromosome was the only chromosome to exhibit such a phenomenon for both sets of DEGs. We then determined if this skew in DEGs translated into overall reduced X chromosome expression compared to autosomes. We found that there was significantly lower expression of X chromosome genes versus autosomes, when comparing the log_2_ fold change between *Taf4b*-heterozygous and *Taf4b*-deficient oocytes (**Fig 3C**). Two similar but slightly different dosage compensation calculation methods, including X:A ratio and relative X expression (RXE), further support that the expression of X chromosome genes is reduced in E16.5 *Taf4b*-deficient oocytes (outliers not plotted) (**Fig 3D-E**). However, we did not see a significant difference in X chromosome expression when looking at E14.5 oocytes (**Fig S4C-E**).

**Figure 3.**
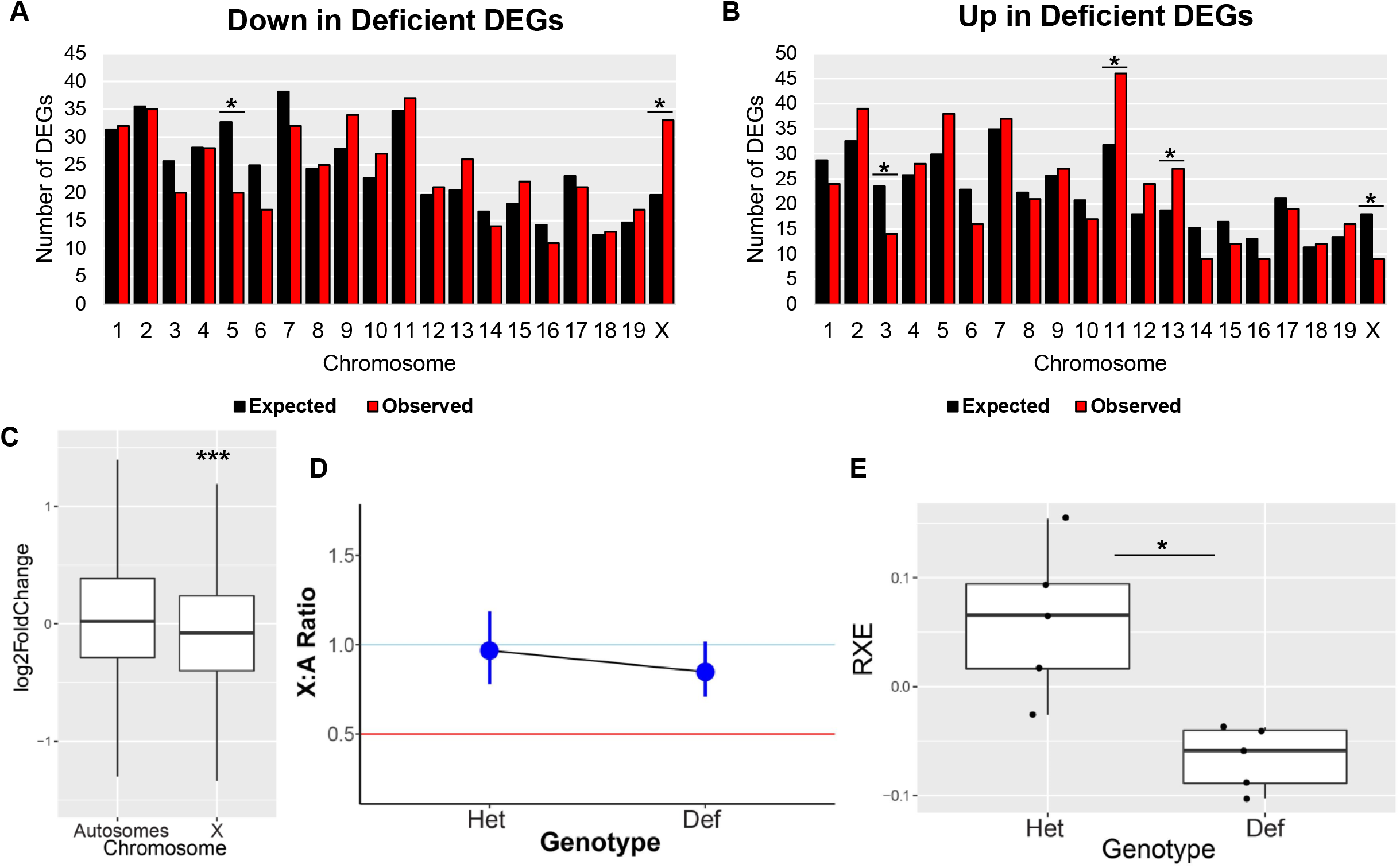
X chromosome gene expression in E16.5 **Taf4b**-deficient oocytes. (A-B) Graphs of expected (black bars) and observed (red bars) numbers of DEGs on each chromosome for the “down in deficient” (A) and “up in deficient” (B) DEGs. * = p < 0.05, chi-square test. (C) Boxplots of log_2_ fold change values from DESeq2 for genes on autosomes versus the X chromosome (outliers removed). *** = p < 0.0001, Welch’s t-test. (D) X:A ratio plot calculated through pairwiseCI after filtering for average TPM > 1 comparing Het X:A ratio to **Taf4b**-deficient X:A Ratio. (E) Boxplots of relative X expression (RXE) calculations after filtering for average TPM > 1 and adding pseudocounts for log-transformation for **Taf4b**-heterozygous and -deficient samples, * = p < 0.05, Welch’s t-test.

Ohno’s hypothesis postulates that the expression of the X chromosome is uniquely regulated so that “housekeeping genes” on the X largely remain on par with autosomal housekeeping gene expression. In Sangrithi et al., 2017, the authors annotated the genome for genes expressed (FPKM ≥1) in all tissues they sampled (Sangrithi et al. 2017). We used this set of ubiquitously expressed genes to see if the effects of *Taf4b*-deficiency on X chromosome expression were specific to ubiquitously expressed genes. We found that 39% of our DEGs were members of the ubiquitous genes list (**Fig S5A**), which is higher than the 25% of all genes being ubiquitously expressed. However, when we plotted the log_2_ fold change of ubiquitous genes on the X chromosome and autosomes, there was no significant difference between these populations (**Fig S5B, Table S6**). Taken together, these data indicate that *Taf4b*-deficiency affects the expression of the X chromosome but it is unclear if *Taf4b* plays a direct role in dosage compensation.

### Overlap of mouse genes deregulated by *Taf4b*-deficiency and Turner Syndrome

As this is the first link of *Taf4b* to the regulation of X-linked gene expression, we decided to compare it to a mouse model of Turner Syndrome (TS). TS is a chromosomal disorder where a female individual has one intact X chromosome and a second X chromosome either missing or severely compromised (Gravholt et al. 2019). We re-processed the raw data from Sangrithi et al., 2017, where the researchers used Oct4-EGFP mice covering 4 developmental time points in female XX and XO germ cells, ranging from E9.5 to E18.5 (**Fig 4A**; Sangrithi et al. 2017). *Taf4b* expression was not significantly different between the karyotypes at E9.5 and E14.5, but it was significantly reduced in E15.5 and E18.5 XO oocytes, whereas *Taf4a* expression was not significantly different at any time point (**Fig 4B-C, Table S7**). To examine the potential overlap of transcriptomic effects between TS and *Taf4b*-deficiency, we compared their DEGs. We first compared E15.5 TS DEGs to our E16.5 *Taf4b* DEGs and found 243 genes shared between the two gene sets, was a significant overlap (p < 0.05, hypergeometric test) (**Fig 4D**). When we used these 243 genes as input for GO analysis, we found DNA-related categories enriched such as “DNA repair” and “covalent chromatin modification” (**Fig 4E**). We then compared E18.5 TS DEGs to our E16.5 *Taf4b* DEGs and found 439 genes shared between the two lists, which was also a significant overlap (p < 0.05, hypergeometric test) (**Fig 4F**). When we used these 439 genes as input for GO analysis, we again found DNA-related categories enriched such as “DNA repair” (**Fig 4G**). These data indicate that there are shared transcriptomic effects of both TS and *Taf4b*-deficiency in mouse embryonic oocytes, and that these shared effects are related to functions concerning DNA repair and chromatin modification.

**Figure 4.**
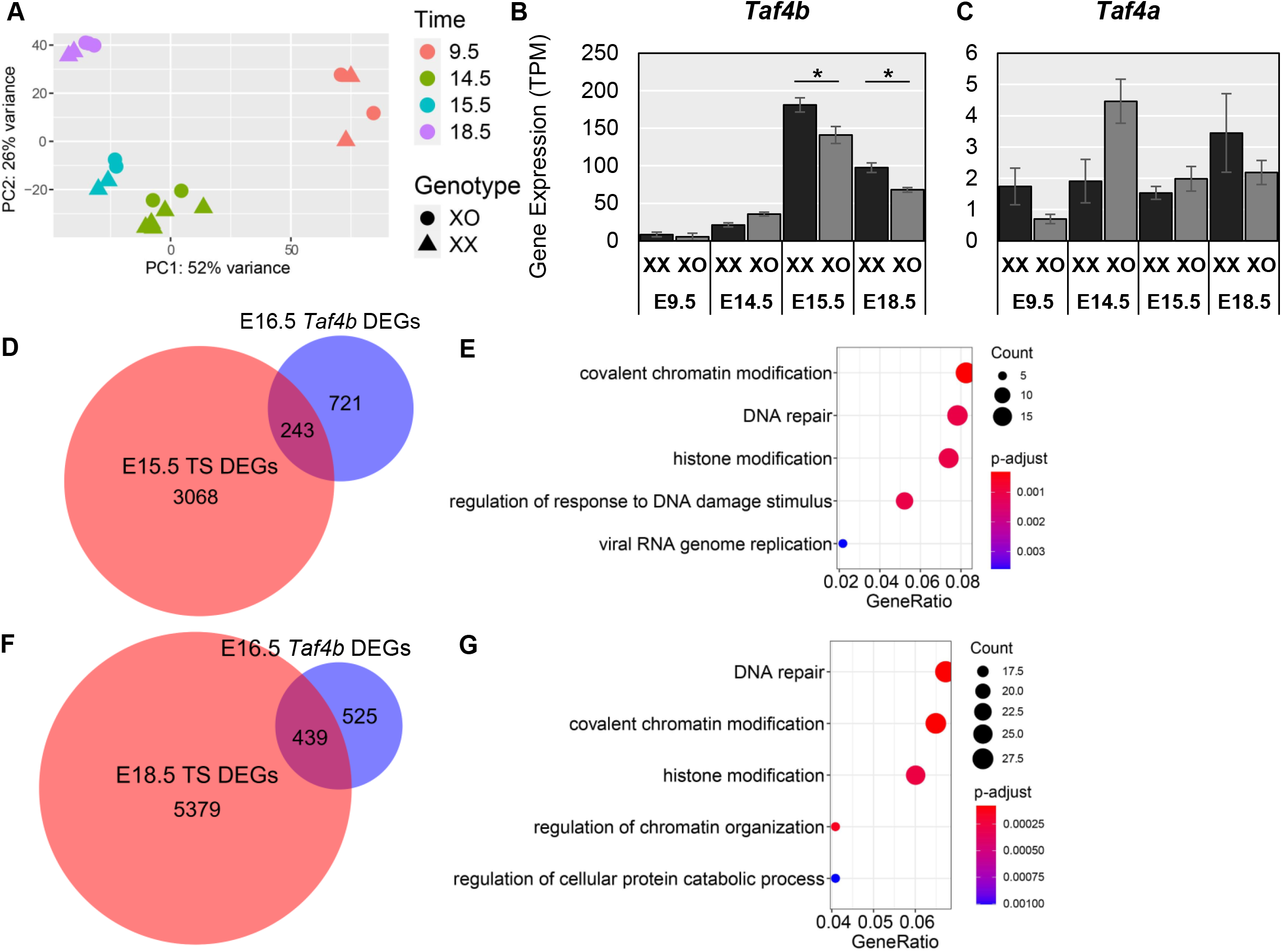
Effects of Turner Syndrome on **Taf4b** and similarities in transcriptomes. (A) PCA plot of the sorted oocytes from Sangrithi et al., 2017, labeled based on embryonic time point and geno type. (B) Expression levels of **Taf4b** in XX versus XO female oocytes (* = p < 0.05, avg TPM > 1). Error bars indicate ± standard error of the mean (SEM). (C) Expression levels of *Taf4a* in XX versus XO female oocytes. Error bars indicate ± standard error of the mean (SEM). (D) Venn diagram of E16.5 **Taf4b** DEG list compared with E15.5 TS DEGs (protein-coding, p-adj < 0.05, avg TPM > 1). Significant overlap in Venn diagram (p < 0.0001, hypergeometric test). (E) GO biological process dotplot for the 243 DEGs shared between E16.5 **Taf4b** and E15.5 TS RNA-seq experiments. (F) Venn diagram of E16.5 **Taf4b** DEG list compared with E18.5 TS DEGs (protein-coding, p-adj < 0.05, avg TPM > 1). Significant overlap in Venn diagram (p < 0.0001, hypergeometric test). (G) GO biological process dotplot for the 439 DEGs shared between E16.5 *Taf4b* and E18.5 TS RNA-seq experiments.

We observed similar results in an independent dataset. By comparing *Taf4b* expression in XX and XO cells that had been differentiated from mouse embryonic stem cells *in vitro* we found that *Taf4b* expression was lower in the XO cells that best resembled late embryonic oocytes (**Table S8**). Interestingly, when the cells had been further differentiated to be similar to early postnatal oocytes, the trend reversed with *Taf4b* expression being significantly higher in mature oocyte-like cells derived from XO cells. This corroborates the reduction in *Taf4b* expression in mature mouse oocytes and suggests that oocyte expression of *Taf4b* normally decreases postnatally. In contrast, significant differences in *Taf4a* expression did not occur until the latter stages of differentiation that best resembled postnatal oocytes (**Fig S7**).

### CUT&RUN identifies putative direct targets of TAF4b in E16.5 germ cells

To understand which DEGs identified in our E16.5 RNA-seq experiment were likely to be direct targets of TAF4b, we performed Cleavage Under Targets and Release Using Nuclease (CUT&RUN), a technique to map binding sites of proteins or histone modification in the genome, in embryonic germ cells. We isolated E16.5 female and male germ cells using FACS and examined the genomic localization of TAF4b, using H3K4me3 as a positive control and to mark promoter regions, and IgG as a negative control. CUT&RUN data analysis using the program Homer identified 11,636 female H3K4me3 peaks, 570 female TAF4b peaks, 59,307 male H3K4me3 peaks, and 3,152 male TAF4b peaks (**Table S9**). We also found that 94% and 73% of TAF4b peaks were classified as localizing to promoters/transcription start site (“Promoter-TSS”) for female and male germ cells, respectively (**Fig 5A**). Of all the TAF4b peaks, 152 overlapped between the sexes (**Fig 5B**). To see if the transcriptomic effects of TAF4b loss on the X chromosome arise from greater X chromosome localization, we plotted the expected versus observed number of peaks using the TAF4b peaks that were categorized as “Promoter-TSS” and found that there were fewer X chromosome peaks than expected in females and no significant difference in males (**Fig 5C-D**). Given that we found that there are more DEGs between *Taf4b*-heterozyous and -deficient oocytes on the X chromosome than expected, this suggests that there might be an indirect but disproportionately high effect of TAF4b on the X chromosome in E16.5 oocytes. However, we cannot claim this experiment thoroughly annotated all TAF4b peaks in the germ cell genome and so future CUT&RUN experiments may reveal more TAF4b localization on the X chromosome.

**Figure 5.**
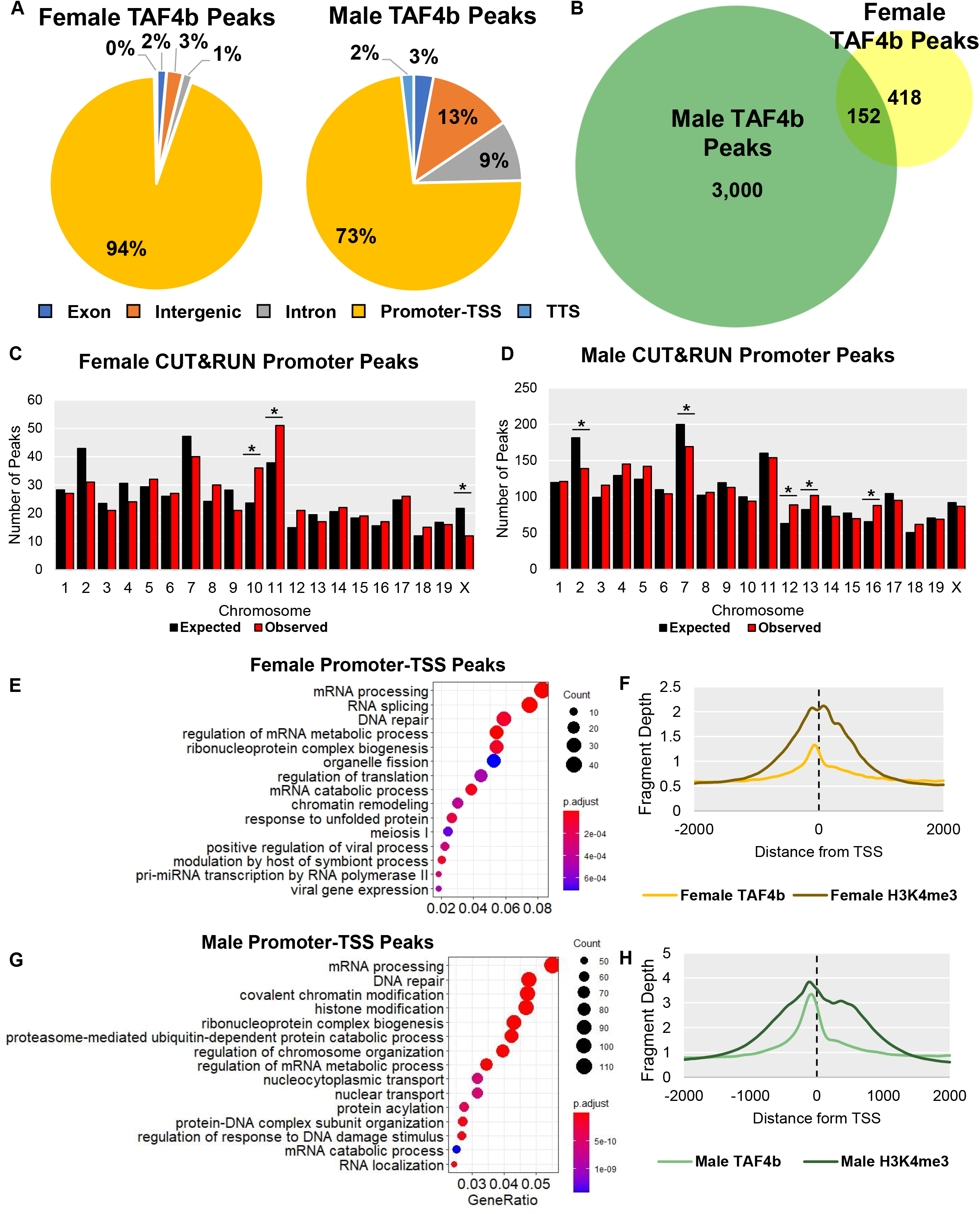
E16.5 germ cell CUT&RUN identifies direct targets of TAF4b. (A) Pie charts of TAF4b peak locations in female and male germ cell CUT&RUN. (B) Venn diagram of peaks shared between female and male germ cell CUT&RUN. Graphs of expected (black bars) and observed (red bars) numbers of promoter peaks on each chromosome for the female (C) and male (D) CUT&RUN peaks (* = p < 0.05, chi-square test). (E) GO biological process dotplot for the female CUT&RUN peaks categorized as “promoter-TSS”. (F) Average enrichment of TAF4b near TSSs (dotted line) in female germ cells. (G) GO biological process dotplot for the male CUT&RUN peaks categorized as “promoter-TSS”. (H) Average enrichment of TAF4b near TSSs (dotted line) in male germ cells.

When we performed GO analysis of TAF4b-bound gene promoters-TSSs, familiar categories appeared for both female and male germ cells. In female germ cells, we found categories related to mRNA processing, translation, and meiosis (**Fig 5E, Table S9**), with multiple genes of interest such as *Mlh1* and *Meioc* (**Fig S8**). When plotting the enrichment profile of TAF4b relative to TSSs, we found the highest TAF4b enrichment upstream of the TSS (**Fig 5F**). In male germ cells, we found categories related to mRNA, DNA repair, and chromatin (**Fig 5G**), with multiple TAF4b peaks of interest such as Smad4 and Bub3 (**Fig S8**). Moreover, we found that in both female and male germ cells TAF4b bound at the promoter of *Dazl*, a gene that controls gamete development, which was previously shown to have TAF4b bound to its promoter by ChIP-qPCR (Grive et al. 2016). (**Fig S8A**). When plotting the enrichment profile of TAF4b relative to TSSs in these cells, we again found the highest TAF4b enrichment upstream of the TSS (**Fig 5H**). However, it is clear when looking at some gene tracks (Ep300, for example) that even when a TAF4b peak is identified in only one of the sexes, there is enrichment of TAF4b in the same location in the other sex, suggesting that some TAF4b binding sites are below the limit of detection by peak calling (**Fig S8**). The CUT&RUN experiment identified shared, and also unique TAF4b binding sites in female and male germ cells, especially around TSSs. This CUT&RUN experiment allowed us to begin inquiring as to which DEGs identified in our RNA-seq experiment were putative direct targets of TAF4b. When comparing our DEGs to the “Promoter-TSS” peaks of TAF4b, we found 202 DEGs that had at least one peak near their TSS (**Fig 6A**). GO analysis of these peaks found that the main categories enriched in these data included DNA repair and translation (**Fig 6B**). As examples of TAF4b-bound DEGs, we present gene tracks for *JunD, Sp1*, and *Fmr1* (**Fig 6C**). *JunD* was upregulated in *Taf4b*-deficient oocytes, which is a component of the AP-1 transcription factor complex (Mechta-Grigoriou et al. 2001). *Sp1* was also upregulated in *Taf4b*-deficient oocytes and is a transcription factor (Vizcaíno et al. 2015). These data suggest that TAF4b likely directly regulates transcription factors and *Fmr1*, a common genetic cause of POI and an upregulated gene in *Taf4b*-deficient oocytes. Furthermore, we identified meiotic genes that were DEGs in and TAF4b-bound, including *Mlh1* (**Table S9**).

**Figure 6.**
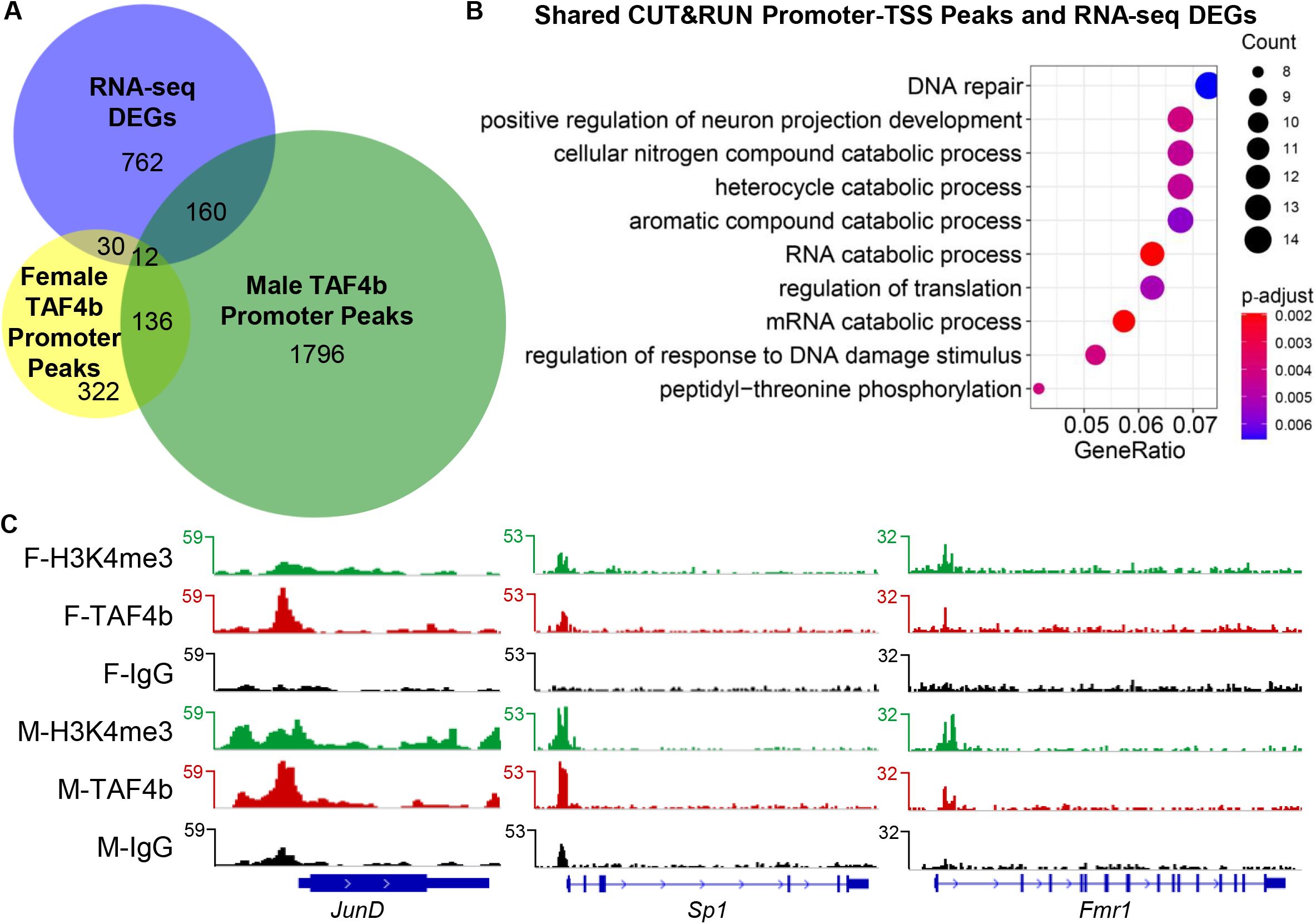
E16.5 CUT&RUN identifies putative direct targets of TAF4b in germ cells. (A) Venn diagram of CUT&RUN promoter peaks and RNA-seq DEGs. (B) Biological process GO dotplot of the 202 genes that are in the list of DEGs and had a promoter-TSS peak in at least one of the two germ cell samples. (C) Gene track of *JunD*, a DEG that had a promoter-TSS peak in both female and male CUT&RUN experiments. (D) Gene track of *Sp1*, a DEG that had a promoter-TSS peak in only the female CUT&RUN. (E) Gene track of *Fmr1*, a DEG that had a promoter-TSS peak in only the male CUT&RUN.

We then identified conserved promoter elements in TAF4b peaks and were surprised to find that TATA-box was not among the top 10 TAF4b motifs in female nor male germ cells (**Fig 7A-B**). Instead, motifs from the Sp/KLF family of transcription factors dominated the lists, especially in the female. When we plotted the average distance of a TAF4b “Promoter-TSS” peak to the nearest TSS, the distance was −88 base pairs for females and −70 base pairs for males (**Fig 7C**), suggesting that TAF4b may be playing a role outside of canonical TFIID in mouse embryonic germ cells. However, more canonical functions of TAF4b cannot be ruled out, as “TATA-Box (TBP)/Promoter” did appear as a significantly enriched motif in both sexes, it was ranked 162 in females and 126 in males (**Table S9**). These data combining RNA-seq and CUT&RUN of TAF4b suggest that TAF4b directly regulates meiotic genes and transcription factors in oocytes, perhaps through a non-canonical protein complex that prioritizes other motifs over the TATA-box (discussed below).

**Figure 7.**
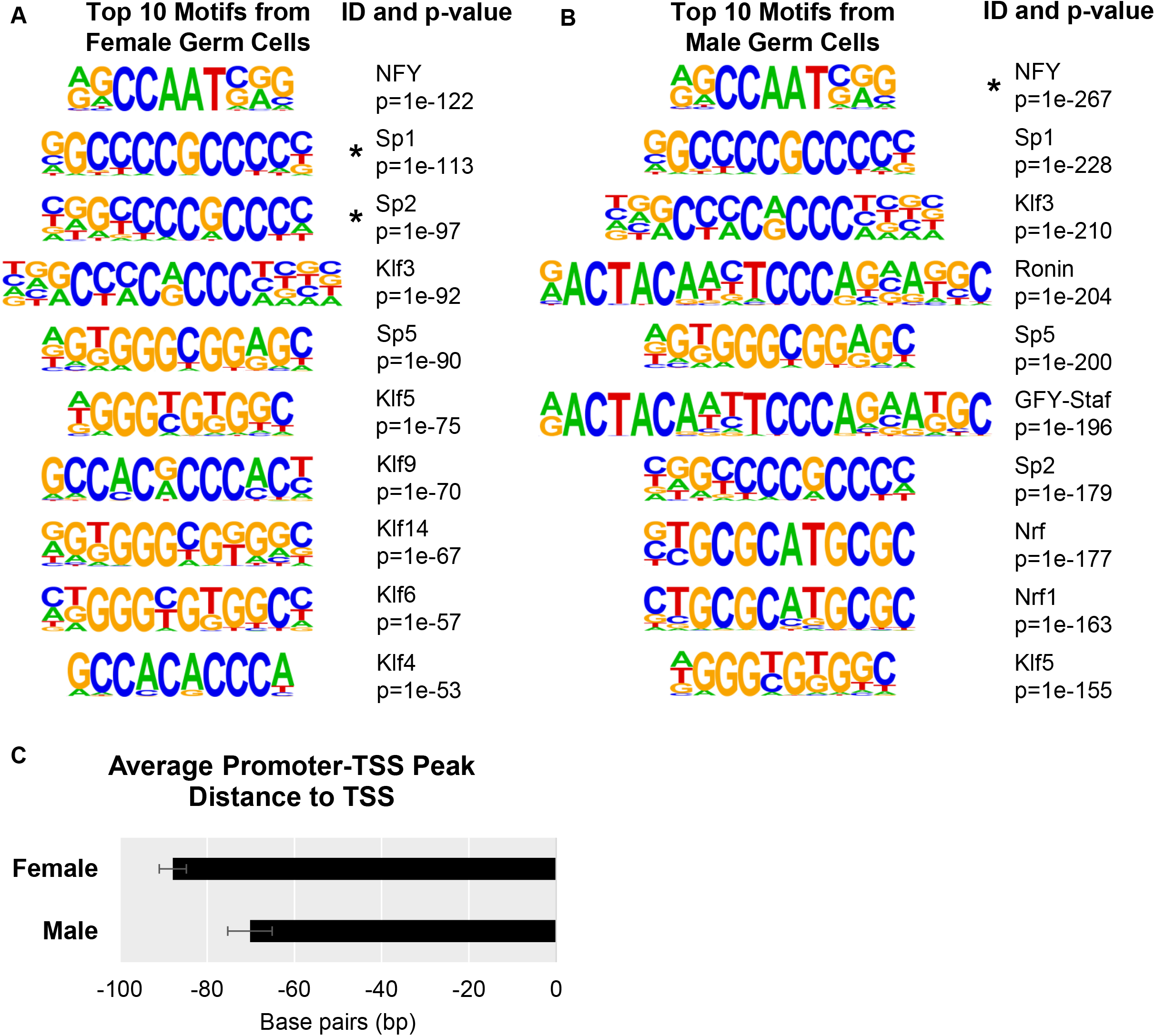
TATA-box not among best candidates for TAF4b motifs. (A) Top ten female motifs enriched at TAF4b peaks, the promoter ID, and the associated p-value. Asterisks indicate that gene was also identified as TAF4b-bound in the CUT&RUN sample. (B) Top ten male motifs enriched at TAF4b peaks, the promoter ID, and the associated p-value. Asterisks indicate that gene was also identified as TAF4b-bound in the CUT&RUN sample. (C) Average distance of TAF4b “promoter-TSS” peak to the associated TSS. Error bars indicate ± standard error of the mean (SEM).

## DISCUSSION

Proper establishment of the ovarian reserve is essential for the reproductive capacity of female mammals, including both humans and mice. This healthy establishment of female gametes is orchestrated through complex oocyte transcription networks that must also properly distinguish germ cell and somatic cell lineages. In addition to the more well-known enhancer-bound transcription activators and repressors, tissue-selective components of the basal transcription machinery can help impart such exquisite regulatory control (Freiman 2009; Goodrich and Tjian 2010). We have previously shown that the TAF4b subunit of the TFIID complex is required for proper establishment of the ovarian reserve in the mouse (Grive et al. 2014, 2016; Lovasco et al. 2010b). However, the network of genes regulated by TAF4b to accomplish this critical task has been elusive, until now. Here we show that TAF4b directly and indirectly regulates genes essential for proper meiotic progression during early oocyte differentiation. Integration of RNA-seq and CUT&RUN data in E16.5 mouse oocytes reveals germ cell-intrinsic regulation by TAF4b in the promoter-proximal regions of DNA repair, chromatin modification, and meiosis genes. Furthermore, we discovered an unexpected link between *Taf4b*-deficiency in the mouse to the proper expression of the mouse X chromosome and similarities to the transcriptome of TS, a well-known cause of POI in women (Sangrithi et al. 2017). Surprisingly, TATA-box motifs were not among the top binding motifs in either female or male shown by CUT&RUN nor was the peak enrichment of TAF4b at the expected location. Together, these molecular insights suggest TAF4b directly regulates genes instrumental in establishing the finite ovarian reserve and that TAF4b may have a non-canonical function in mouse oocytes outside of TFIID or in a non-canonical version of TFIID.

TFIID was first discovered as a large multiprotein complex required for activator-dependent RNA polymerase II transcription (Dynlacht et al. 1991; Reinberg et al. 1987). Characterization of the composition of TFIID revealed a key DNA binding subunit, TBP, that binds directly to the TATA-box found at the −25 nucleotide position in relation to the TSS of many genes (Hoey et al. 1990). In addition, TAF2 and TAF1 were shown to make base-specific contacts with the initiator (INR) and downstream promoter element (DPE), respectively (Smale and Baltimore 1989; Verrijzer et al. 1995). Cryoelectron microscopy structures of TFIID confirms this composite TFIID binding region (−40 to +40) often referred as the core promoter (He et al. 2016). Given this regulatory complexity of the core promoter, TFIID can bind and activate TATA-less promoters likely through these other conserved core promoter elements. Surprisingly, our embryonic germ cell CUT&RUN data for TAF4b centers its peak of binding to G-C and CCAAT-box sequences either at −90 (female) or −70 (male) upstream, but still proximal to the TSS. These sequences are well-known binding sites for specificity protein 1 (SP1) and nuclear factor y (NFY) transcription factors that are known to play extensive roles in promoter proximal transcription. Although we do not yet know the significance of these binding sites and the occupancy of TAF4b, we speculate that their association may bridge critical proximal and core promoter bound elements to drive germ cell-specific programs of gene transcription required for at least this discrete step of oogenic differentiation. Although, we did not identify the enrichment of TAF4b at the TATA, INR, or DPE core promoter elements, it is likely that additional subunits of the TAF4b-containg TFIID complex more closely associate with these sequences. Further molecular investigations are required to sort out the germ cell-specificity of this regulatory logic and the composition of this germ cell-specific version of TFIID.

Similar diversification of selective TFIID subunits has occurred within germline development of highly distant organisms including in insects, vertebrates and plants. In Drosophila, several testis-specific TAFs (tTAFs) play a critical role in regulating transcription and the timing of spermatogenic differentiation and a germ cell expressed TBP paralog, TBP-related factor 2 (TRF2) is required for oogenesis (Gazdag et al. 2009; Hiller et al. 2004). The mouse ortholog of TRF2, called TBPL1, is required for spermiogenesis as is TAF7l that is coordinately expressed with TAF4b in early meiotic oocytes (Zhou et al. 2013a). Interestingly, TBPL1, TAF7l, and TAF9b have also been shown to be critical for muscle and adipocyte differentiation, indicating that this diversification of TFIID subunits has evolved to regulate both somatic and germ cell differentiation, sometimes via the identical subunit (Herrera et al. 2014; Zhou et al. 2013b). The most striking parallel of the early meiotic transcription and chromatin functions of TAF4b shown here lie with a natural variant of TAF4b found in Arabidopsis (AtTAF4b) (Lawrence et al. 2019). This recent study has identified a similar timing of the meiocyte transcriptome regulated by AtTAF4b as we show here for mouse TAF4b. Although arising independently in the plant and animal kingdoms, there appears to be some common transcription and/or chromatin state that is regulated by TAF4b to ensure the fidelity of meiotic recombination and early oogenesis. We have previously found other TFIID subunits such as *Taf7l* and *Taf9b* to be preferentially and dynamically expressed in embryonic mouse oocytes (Gura et al. 2020). One hypothesis to explain this data is that a germ cell-specific version of TFIID may exhibit different characteristics and targets than canonical TFIID.

In addition to illuminating the molecular underpinnings of TAF4b function, we discovered unexpected overlaps between *Taf4b*-deficiency and other known causes of POI. Many TS individuals experience POI, which includes not reaching or delayed menarche and primary amenorrhea. Recent research has suggested that excessive prenatal oocyte loss may underlie the ovarian insufficiency in TS. Excessive oocyte attrition at the perinatal DNA damage checkpoint, where oocytes that have not resolved DNA damage or still contain asynapsed chromosomes are eliminated, results in a depleted ovarian reserve and its downstream consequences. Observing extensive asynapsis and deregulation of the X chromosome in *Taf4b*-deficient E16.5 oocytes prompted us to compare *Taf4b*-deficiency and a mouse model of TS where females contain a single X chromosome (XO) (Grive et al. 2016). The extensive overlap of the deregulated gene expression in XO and *Taf4b*-deficient early meiotic oocytes was striking. Importantly, *Taf4b* expression itself was compromised in the XO oocytes, indicative of potential mutual regulation between these two genetic changes. A recent report uncovered a unique mechanism of XX dosage compensation in human primordial oocytes and it is possible that TAF4b plays an integral role in this sexually dimorphic mechanism of X-chromosome regulation (Chitiashvili et al. 2020). Another interesting parallel is the association of X chromosome-encoded *Fmr1* gene with TS and *Taf4b*-deficiency. *Taf4b* expression is significantly correlated with *Fmr1* in embryonic human ovaries, mutation of *Fmr1* is one the most common underlying genetic causes of POI, and TS individuals are missing one copy of Fmr1. Here we show that TAF4b directly associates with the proximal promoter region of *Fmr1* and the loss of TAF4b increases its mRNA abundance. Given the clear links between *Taf4b*-deficiency, TS, FXPOI, and POI presented here, we suspect that a core of the genes identified in this study are required for the proper development of the ovarian reserve in humans and if that quorum is not reached a similar cascade of dysregulated gene expression occurs. Understanding these and other causes of POI will clarify the best ways to manage these related infertility syndromes and improve assisted reproduction therapies.

## MATERIALS AND METHODS

### Ethics statement

This study was approved by Brown University IACUC protocol #21-02-0005. The primary method of euthanasia is CO_2_ inhalation and the secondary method used is cervical dislocation both as per American Veterinary Medical Association (AVMA) guidelines on euthanasia.

### Mice

Mice that were homozygous for an *Oct4-EGFP* transgene (The Jackson Laboratory: B6;129S4-*Pou5f1^tm2Jae^*/J) were mated for CUT&RUN collections. Mice that were homozygous for an *Oct4-EGFP* transgene (The Jackson Laboratory: B6;129S4-*Pou5f1^tm2Jae^*/J) and heterozygous for the *Taf4b*-deficiency mutation (in exon 12 of the 15 total exons of the *Taf4b* gene that disrupts the endogenous *Taf4b* gene) were mated for mRNA collections. Timed matings were estimated to begin at day 0.5 by evidence of a copulatory plug. The sex of the embryos was identified by confirming the presence or absence of testicular cords. Genomic DNA from tails was isolated using Qiagen DNeasy Blood & Tissue Kits (Cat #: 69506) for PCR genotyping assays.

All animal protocols were reviewed and approved by Brown University Institutional Animal Care and Use Committee and were performed in accordance with the National Institutes of Health Guide for the Care and Use of Laboratory Animals. Gonads were dissected out of embryos into cold PBS.

### Embryonic gonad dissociation and fluorescence-activated cell sorting

To dissociate gonadal tissue into a single-cell suspension, embryonic gonads were harvested and placed in 0.25% Trypsin/EDTA and incubated at 37°C for 15 and 25 minutes for E14.5 and E16.5 ovaries, respectively, as previously described (Gura et al. 2020). Eppendorf tubes were flicked to dissociate tissue halfway through and again at the end of the incubation. Trypsin was neutralized with FBS. Cells were pelleted at 1,500 RPM for 5 minutes, the supernatant was removed, and cells were resuspended in 100 μL PBS. The cell suspension was strained through a 35 μm mesh cap into a FACS tube (Gibco REF # 352235). Propidium iodide (1:500) was added to the cell suspension as a live/dead distinguishing stain. Fluorescence-activated cell sorting (FACS) was performed using a Becton Dickinson FACSAria III in the Flow Cytometry and Cell Sorting Core Facility at Brown University. A negative control of a non-transgenic mouse gonad was used for each experiment to establish an appropriate GFP signal baseline. Dead cells were discarded and the remaining cells were sorted into GFP^+^ and GFP^-^ samples in PBS at 4°C for each embryo.

For RNA-seq analysis, GFP^+^ cells from each individual embryo were kept in separate tubes and were then spun down at 1,500 RPM for 5 minutes, had PBS removed, and were then resuspended in Trizol (ThermoFisher # 1556026). If samples had roughly less than 50 µL of PBS in the tube, Trizol was added immediately. The number of cells for each sample can be found in **Table S10**. Samples were stored at −80°C.

For CUT&RUN germ cells from all the gonads for each sex were pooled prior to FACS. Sorted cells were then spun down at 1,500 RPM for 5 minutes and were resuspended in 300 µL of PBS, then split into three Eppendorf tubes. These three tubes of germ cells were then used for CUT&RUN. The number of cells for each sample were as follows: female germ cell samples had 42,416 cells per tube (obtained from 12 embryos) and male germ cell samples had 55,829 cells per tube (obtained from 22 embryos).

### Single cell RNA-seq data analysis

All computational scripts regarding single cell RNA-seq (scRNA-seq) used in this publication are available to the public: https://github.com/mg859337/Gura_et_al._2021/tree/main/scRNA-seq_data_analysis. SRP193506 and SRP188873 were downloaded from NCBI SRA onto Brown University’s high-performance computing cluster at the Center for Computation and Visualization. The fastq files were aligned using Cell Ranger (v 5.0.0) count and then aggregated using Cell Ranger aggr. The resulting output from aggr was used as input for Seurat (v 3.9.9) in RStudio (R v 4.0.2) (Stuart et al. 2019). Seurat was used to select for *Dazl*-positive (*Dazl* > 0), high-quality (nFeature_RNA > 1000, nFeature_RNA < 5000, nCount_RNA > 2500, nCount < 30000, percent mitochondrial genes < 5%) oocytes. This data was then passed to Monocle3 (v 0.2.3) for pseudotime analysis and generating UMAP and gene expression (Cao et al. 2019; Qiu et al. 2017; Trapnell et al. 2014). The cloupe file created from Cell Ranger aggr was used as input for Loupe Cell Browser (v 5.0), where the same filtering steps were used (*Dazl* > 0, Feature Threshold > 1000, Feature Threshold < 5000, UMI Threshold > 2500, UMI Threshold < 30000, Mitochondrial UMIs < 5%). These filtered cells were then split into *Taf4b*-on (*Taf4b* > 0) and *Taf4b* -off (Taf4b = 0) and then “Locally Distinguishing” was run for Significant Feature Comparison. The list of genes significantly associated with *Taf4b*-on cells (**Table S1**) was used as input for ClusterProfiler (v 3.16.1) to create a dotplot of significantly enriched gene ontology (GO) categories (Yu et al. 2012).

### RNA-sequencing

Embryonic germ cells resuspended in Trizol were shipped to GENEWIZ (GENEWIZ Inc., NJ) on dry ice. Sample RNA extraction, sample QC, library preparation, sequencing, and initial bioinformatics were done at GENEWIZ. RNA was extracted following the Trizol Reagent User Guide (Thermo Fisher Scientific). Glycogen was added (1 µL, 10 mg/mL) to the supernatant to increase RNA recovery. RNA was quantified using Qubit 2.0 Fluorometer (Life Technologies, Carlsbad, CA, USA) and RNA integrity was checked with TapeStation (Agilent Technologies, Palo Alto, CA, USA) to see if the concentration met the requirements.

SMART-Seq v4 Ultra Low Input Kit for Sequencing was used for full-length cDNA synthesis and amplification (Clontech, Mountain View, CA), and Illumina Nextera XT library was used for sequencing library preparation. The sequencing libraries were multiplexed and clustered on a lane of a flowcell. After clustering, the flowcell was loaded onto an Illumina HiSeq 4000 according to manufacturer’s instructions. The samples were sequenced using a 2×150 Paired End (PE) configuration. Image analysis and base calling were conducted by the HiSeq Control Software (HCS) on the HiSeq instrument. Raw sequence data (.bcl files) generated from Illumina HiSeq were converted into fastq files and de-multiplexed using bcl2fastq (v. 2.17). One mismatch was allowed for index sequence identification.

### RNA-seq data analysis

All computational scripts regarding RNA-seq used in this publication are available to the public: https://github.com/mg859337/Gura_et_al._2021/tree/main/RNA-seq_data_analysis. Datasets SRP059601 and SRP059599 were from NCBI SRA. All raw fastq files were initially processed on Brown University’s high-performance computing cluster. Reads were quality-trimmed and had adapters removed using Trim Galore! (v 0.5.0) with the parameters –nextera -q 10. Samples before and after trimming were analyzed using FastQC (v 0.11.5) for quality and then aligned to the Ensemble GRCm38 using HiSat2 (v 2.1.0) (Andrews 2010; Pertea et al. 2016). Resulting sam files were converted to bam files using Samtools (v 1.9) (Li et al. 2009). E14.5 heterozygous bamfiles were downsampled because these samples had been sequenced more deeply than their wild-type and deficient counterparts.

To obtain TPMs for each sample, StringTie (v 1.3.3b) was used with the optional parameters -A and -e. A gtf file for each sample was downloaded and, using RStudio (R v 4.0.2), TPMs of all samples were aggregated into one comma separated (csv) file using a custom R script. To create interactive Microsoft Excel files for exploring the TPMs of each dataset: the csv of aggregated TPMs was saved as an Excel spreadsheet, colored tabs were added to set up different comparisons, and a flexible Excel function was created to adjust to gene name inputs. To explore the Excel files, please find the appropriate tab (named “Quick_Calc”) and type in the gene name of interest into the highlighted yellow boxes. There is an Excel file for each dataset analyzed as a supplementary table.

To obtain count tables, featurecounts (Subread v 1.6.2) was used (Liao et al. 2014). Metadata files for dataset were created manually in Excel and saved as a csv. These count tables were used to create PCA plots by variance-stabilizing transformation (vst) of the data in DESeq2 (v 1.22.2) and plotting by ggplot2 (v 3.1.0) (Love et al. 2014; Wickham 2016). DESeq2 was also used for differential gene expression analysis, where count tables and metadata files were used as input. We accounted for the litter effect in our mouse oocytes by setting it as a batch parameter in DESeq2. For the volcano plot, the output of DESeq2 was used and plotted using ggplot2. DEG lists were used for ClusterProfiler (v 3.16.1) input to create dotplots of significantly enriched gene ontology (GO) categories for all DEGs, Down in Def DEGs, and Up in Def DEGs. Physical, highest-confidence protein-protein interactions were identified using STRING, with unconnected proteins not shown in the image (Szklarczyk et al. 2019).

For X chromosome analysis, expected numbers of Down in Def and Up in Def DEGs per chromosome were calculated by dividing the average number of observations per chromosome by the average number of total genes per chromosome. Chi-square values and p-values were calculated using the GraphPad QuickCalcs chi-square function, where observed and expected frequencies were used as input (https://www.graphpad.com/quickcalcs/chisquared1/, accessed Jan 2021). Boxplots of log2 fold change between the autosomes and X chromosomes used the output of DESeq2 as input, based on other publications comparing autosomal and X chromosome expression (Hirota et al. 2018). The X:A ratio was calculated using pairwiseCI (v. 0.1.27), a bootstrapping R package, after filtering genes for an average TPM > 1 (Duan et al. 2019b; Sangrithi et al. 2017). The RXE was calculated using a custom R script based after filtering genes for an average TPM > 1 and adding pseudocounts for log transformation (log2(x+1)), based on other RXE publications (Duan et al. 2019a; Jue et al. 2013). The “ubiquitous genes” from Sangrithi et al., 2017 were converted from gene names to Ensembl IDs, first by using ShinyGO to convert IDs through the “Genes” tab (Ge et al. 2020). Genes that were not mapped were then used as input for DAVID gene ID conversion, any remaining unconverted gene names were manually entered into the Ensembl database to find matches (Howe et al. 2021; Huang et al. 2007). Venn diagrams were created using BioVenn (Hulsen et al., 2008). All plots produced in RStudio were saved as an EPS file type and then opened in Adobe Illustrator in order to export a high-quality JPEG image.

### CUT&RUN

The CUT&RUN performed in E16.5 germ cells followed the protocol in Hainer and Fazzio, 2019. CUT&RUN antibodies were as follows: polyclonal rabbit TAF4b (as previously described (Grive et al. 2016)), monoclonal rabbit H3K4me3 (EMD Millipore # 05-745R), rabbit IgG (ThermoFisher # 02-6102), pA-MNase was a generous gift from Dr. Thomas Fazzio.

For library preparation, the KAPA HyperPrep kit (Roche Cat. No 07962363001) was used with New England Biolabs NEBNext Multiplex Oligos for Illumina (NEB #E7335). After library amplification through PCR, libraries were size selected through gel extraction (∼150-650 bp) and cleaned up using the Qiagen QIAquick Gel Extraction Kit (Cat. # 28704). CUT&RUN libraries in EB buffer were shipped to GENEWIZ (GENEWIZ Inc., NJ) on dry ice. Sample QC, sequencing, and initial bioinformatics were done at GENEWIZ.

The sequencing libraries were validated on the Agilent TapeStation (Agilent Technologies, Palo Alto, CA, USA), and quantified by using Qubit 2.0 Fluorometer (Invitrogen, Carlsbad, CA) as well as by quantitative PCR (KAPA Biosystems, Wilmington, MA, USA). The sequencing libraries were clustered on flowcells. After clustering, the flowcells were loaded on to the Illumina HiSeq instrument (4000 or equivalent) according to manufacturer’s instructions. The samples were sequenced using a 2×150bp Paired End (PE) configuration. Raw sequence data (.bcl files) generated from Illumina HiSeq were converted into fastq files and de-multiplexed using bcl2fastq (v. 2.20). One mismatch was allowed for index sequence identification.

### CUT&RUN data analysis

All computational scripts regarding CUT&RUN data analysis used in this publication are available at: https://github.com/mg859337/Gura_et_al._2021/tree/main/CUT%26RUN_data_analysis and based on other CUT&RUN publications (Hainer and Fazzio 2019). All raw fastq files were initially processed on Brown University’s high-performance computing cluster. Reads were quality-trimmed and had adapters removed using Trim Galore! (v 0.5.0) with the parameter -q 10 (https://www.bioinformatics.babraham.ac.uk/projects/trim_galore/). Samples before and after trimming were analyzed using FastQC (v 0.11.5) for quality and then aligned to the Ensemble GRCm39 using Bowtie2 (v 2.3.0). Fastq screen (v 0.13.0) was used to determine the percentage of reads uniquely mapped to the mouse genome in comparison to other species. Resulting sam files were converted to bam files, then unmapped, duplicated reads, and low quality mapped were removed using Samtools (v1.9). Resulting bam files were split into size classes using a Unix script. For calling peaks, merging peaks, and identifying coverage around TSSs, Homer (v 4.10) was used (Heinz et al. 2010). For gene track visualization, the final bam file before splitting into size classes was used as input to Integrative Genomics Viewer (IGV) (Robinson et al. 2011). A custom genome was created using a genome fasta and gtf file for Ensembl GRCm39.

Pie charts were created using data from Homer output and Venn diagrams were created using BioVenn. For X chromosome analysis, expected numbers of promoter peaks per chromosome were calculated by dividing the average number of observations per chromosome by the average number of total protein-coding genes per chromosome. Chi-square values and p-values were calculated using the GraphPad QuickCalcs chi-square function, where observed and expected frequencies were used as input (https://www.graphpad.com/quickcalcs/chisquared1/, accessed Apr 2021). Dotplots of Promoter-TSS peaks were made using ClusterProfiler. To find what TAF4b peaks were shared between female and male germ cells, Homer’s merge function was used. TSS plots were created using the “tss” function of Homer and plotted using Microsoft Excel. All plots produced in RStudio were saved as an EPS file type and then opened in Adobe Illustrator in order to export a high-quality JPEG image.

## ACKNOWLEDGMENTS

We thank Drs. Ashley Webb, Erica Larschan, Mark Johnson, and Kathryn Grive for their helpful input throughout these studies. We thank the Center for Computation and Visualization at Brown University for computational resources for scRNA-seq, RNA-seq, and CUT&RUN data analysis. We thank Kevin Carlson and the Brown University Flow Cytometry and Sorting Facility for expertise completing the flow sorting. The Brown University Flow Cytometry and Sorting Facility has received generous support in part by the National Institutes of Health (NCRR Grant No. 1S10RR021051) and the Division of Biology and Medicine, Brown University. As much of our insights were gained by reprocessing publicly available RNA-seq datasets, we greatly appreciate both the researchers that generated and shared the data initially and the respective repositories for making them available. We are grateful to the NICHD/NIH for their generous support through awards 1F31HD097933, 1F31HD105340, and 1R01HD091848 to MAG, KMA and RNF, respectively and thank the BSF for their generous support.

## Author Contributions

MAG, conception and design of all experiments, collection and assembly of data, data analysis and interpretation, manuscript writing; SR, collection and assembly of data, data analysis and interpretation; KMA, collection and assembly of data, data analysis and interpretation; KAS, mouse colony management, collection of ovarian tissue and cells, data analysis and interpretation; TW, reagents and technical expertise for CUT&RUN data collection, analysis, experimental design, and data interpretation; HK, technical expertise and data interpretation; JMAT, technical expertise and data interpretation; TGF, reagents and technical expertise for CUT&RUN data collection and analysis, experimental design and data interpretation; RNF, conception and design of all experiments, data interpretation, manuscript writing and financial support.

## REFERENCES

Andrews S. 2010. FastQC: A quality control tool for high throughput sequence data. http://www.bioinformaticsbabrahamacuk/projects/fastqc/ http://www.bioinformatics.babraham.ac.uk/projects/.

Antonova S V., Boeren J, Timmers HTM, Snel B. 2019. Epigenetics and transcription regulation during eukaryotic diversification: the saga of TFIID. Genes Dev 33: 1–15. http://genesdev.cshlp.org/lookup/doi/10.1101/gad.300475.117.

Cao J, Spielmann M, Qiu X, Huang X, Ibrahim DM, Hill AJ, Zhang F, Mundlos S, Christiansen L, Steemers FJ, et al. 2019. The single-cell transcriptional landscape of mammalian organogenesis. Nature 566: 496–502. http://dx.doi.org/10.1038/s41586-019-0969-x.

Chandra A, Copen CE, Stephen EH. 2013. Infertility and impaired fecundity in the United States, 1982-2010: data from the National Survey of Family Growth. Natl Health Stat Report.

Chitiashvili T, Dror I, Kim R, Hsu FM, Chaudhari R, Pandolfi E, Chen D, Liebscher S, Schenke-Layland K, Plath K, et al. 2020. Female human primordial germ cells display X-chromosome dosage compensation despite the absence of X-inactivation. Nat Cell Biol 22: 1436–1446. http://dx.doi.org/10.1038/s41556-020-00607-4.

Duan J, Flock K, Jue N, Zhang M, Jones A, Al Seesi S, Mandoiu I, Pillai S, Hoffman M, O’Neill R, et al. 2019a. Dosage compensation and gene expression of the X Chromosome in sheep. G3 Genes, Genomes, Genet 9: 305–314.

Duan J, Shi W, Jue NK, Jiang Z, Kuo L, O’Neill R, Wolf E, Dong H, Zheng X, Chen J, et al. 2019b. Dosage compensation of the X chromosomes in bovine germline, early embryos, and somatic tissues. Genome Biol Evol 11: 242–252.

Dynlacht BD, Hoey T, Tjian R. 1991. Isolation of coactivators associated with the TATA-binding protein that mediate transcriptional activation. Cell 66: 563–576.

Falender AE, Shimada M, Lo YK, Richards JS. 2005. TAF4b, a TBP associated factor, is required for oocyte development and function. Dev Biol 288: 405–419.

Feng CW, Bowles J, Koopman P. 2014. Control of mammalian germ cell entry into meiosis. Mol Cell Endocrinol 382: 488–497.

Fink DA, Nelson LM, Pyeritz R, Johnson J, Sherman SL, Cohen Y, Elizur SE. 2018. Fragile X Associated Primary Ovarian Insufficiency (FXPOI): Case Report and Literature Review. Front Genet 9: 1–12.

Freiman RN. 2009. Specific variants of general transcription factors regulate germ cell development in diverse organisms. Biochim Biophys Acta - Gene Regul Mech 1789: 161– 166.

Gazdag E, Santenard A, Ziegler-Birling C, Altobelli G, Poch O, Tora L, Torres-Padilla ME. 2009. TBP2 is essential for germ cell development by regulating transcription and chromatin condensation in the oocyte. Genes Dev 23: 2210–2223.

Ge SX, Jung D, Yao R. 2020. ShinyGO: A graphical enrichment tool for ani-mals and plants. Bioinformatics 36: 2628–2629.

Ge W, Wang JJ, Zhang RQ, Tan SJ, Zhang FL, Liu WX, Li L, Sun XF, Cheng SF, Dyce PW, et al. 2021. Dissecting the initiation of female meiosis in the mouse at single-cell resolution. Cell Mol Life Sci 78: 695–713. https://doi.org/10.1007/s00018-020-03533-8.

Goodrich JA, Tjian R. 2010. Unexpected roles for core promoter recognition factors in cell-type-specific transcription and gene regulation. Nat Rev Genet 11: 549–558.

Gravholt CH, Brun S, Andersen NH. 2019. Turner syndrome: mechanisms and management. Nat Rev Endocrinol. http://dx.doi.org/10.1038/s41574-019-0224-4.

Grive KJ, Gustafson EA, Seymour KA, Baddoo M, Schorl C, Golnoski K, Rajkovic A, Brodsky AS, Freiman RN. 2016. TAF4b Regulates Oocyte-Specific Genes Essential for Meiosis. PLoS Genet 12: 1–18.

Grive KJ, Seymour K a., Mehta R, Freiman RN. 2014. TAF4b promotes mouse primordial follicle assembly and oocyte survival. Dev Biol 392: 42–51. http://dx.doi.org/10.1016/j.ydbio.2014.05.001.

Gura MA, Freiman RN. 2018. Primordial Follicle. Encycl Reprod 65–71. https://linkinghub.elsevier.com/retrieve/pii/B9780128012383643945.

Gura MA, Mikedis MM, Seymour KA, De Rooij DG, Page DC, Freiman RN. 2020. Dynamic and regulated TAF gene expression during mouse embryonic germ cell development. PLoS Genet 16: 1–28.

Hainer SJ, Fazzio TG. 2019. High-Resolution Chromatin Profiling Using CUT&RUN. Curr Protoc Mol Biol 126: 1–22.

He Y, Yan C, Fang J, Inouye C, Tjian R, Ivanov I, Nogales E. 2016. Near-atomic resolution visualization of human transcription promoter opening. Nature 533: 359–365. http://dx.doi.org/10.1038/nature17970.

Heinz S, Benner C, Spann N, Bertolino E, Lin YC, Laslo P, Cheng JX, Murre C, Singh H, Glass CK. 2010. Simple Combinations of Lineage-Determining Transcription Factors Prime cis-Regulatory Elements Required for Macrophage and B Cell Identities. Mol Cell 38: 576– 589. http://dx.doi.org/10.1016/j.molcel.2010.05.004.

Herrera FJ, Yamaguchi T, Roelink H, Tjian R. 2014. Core promoter factor TAF9B regulates neuronal gene expression. Elife.

Hiller M, Chen X, Pringle MJ, Suchorolski M, Sancak Y, Viswanathan S, Bolival B, Lin T-Y, Marino S, Fuller MT. 2004. Testis-specific TAF homologs collaborate to control a tissue-specific transcription program. Development 131: 5297–5308.

Hirota T, Blakeley P, Sangrithi MN, Mahadevaiah SK, Encheva V, Snijders AP, ElInati E, Ojarikre OA, de Rooij DG, Niakan KK, et al. 2018. SETDB1 Links the Meiotic DNA Damage Response to Sex Chromosome Silencing in Mice. Dev Cell 47: 645–659.e6.

Hoey T, Dynlacht BD, Peterson MG, Pugh BF, Tjian R. 1990. Isolation and characterization of the Drosophila gene encoding the TATA box binding protein, TFIID. Cell 61: 1179–1186.

Howe KL, Achuthan P, Allen J, Allen J, Alvarez-Jarreta J, Ridwan Amode M, Armean IM, Azov AG, Bennett R, Bhai J, et al. 2021. Ensembl 2021. Nucleic Acids Res 49: D884–D891.

Huang DW, Sherman BT, Tan Q, Kir J, Liu D, Bryant D, Guo Y, Stephens R, Baseler MW, Lane HC, et al. 2007. DAVID Bioinformatics Resources: Expanded annotation database and novel algorithms to better extract biology from large gene lists. Nucleic Acids Res 35: 169–175.

Jue NK, Murphy MB, Kasowitz SD, Qureshi SM, Obergfell CJ, Elsisi S, Foley RJ, O’Neill RJ, O’Neill MJ. 2013. Determination of dosage compensation of the mammalian X chromosome by RNA-seq is dependent on analytical approach. BMC Genomics 14.

Lawrence EJ, Gao H, Tock AJ, Lambing C, Blackwell AR, Feng X, Henderson IR. 2019. Natural Variation in TBP-ASSOCIATED FACTOR 4b Controls Meiotic Crossover and Germline Transcription in Arabidopsis. Curr Biol.

Li H, Handsaker B, Wysoker A, Fennell T, Ruan J, Homer N, Marth G, Abecasis G, Durbin R. 2009. The Sequence Alignment/Map format and SAMtools. Bioinformatics 25: 2078–2079.

Liao Y, Smyth GK, Shi W. 2014. FeatureCounts: An efficient general purpose program for assigning sequence reads to genomic features. Bioinformatics 30: 923–930.

Lovasco L a, Seymour K a, Zafra K, O’Brien CW, Schorl C, Freiman RN. 2010a. Accelerated ovarian aging in the absence of the transcription regulator TAF4B in mice. Biol Reprod 82: 23–34.

Lovasco LA, Gustafson EA, Seymour KA, De Rooij DG, Freiman RN. 2015. TAF4b is required for mouse spermatogonial stem cell development. Stem Cells 33: 1267–1276.

Lovasco LA, Seymour KA, Zafra K, O’Brien CW, Schorl C, Freiman RN. 2010b. Accelerated ovarian aging in the absence of the transcription regulator TAF4B in mice. Biol Reprod 82: 23–34.

Love MI, Huber W, Anders S. 2014. Moderated estimation of fold change and dispersion for RNA-seq data with DESeq2. Genome Biol 15: 550. http://genomebiology.biomedcentral.com/articles/10.1186/s13059-014-0550-8.

Mechta-Grigoriou F, Gerald D, Yaniv M. 2001. The mammalian Jun proteins: Redundancy and specificity. Oncogene 20: 2378–2389.

Pertea M, Kim D, Pertea GM, Leek JT, Salzberg SL. 2016. Transcript-level expression analysis of RNA-seq experiments with HISAT, StringTie and Ballgown. Nat Protoc.

Qiu X, Mao Q, Tang Y, Wang L, Chawla R, Pliner HA, Trapnell C. 2017. Reversed graph embedding resolves complex single-cell trajectories. Nat Methods 14: 979–982.

Reinberg D, Horikoshi M, Roeder RG. 1987. Factors involved in specific transcription in mammalian RNA polymerase II. Functional analysis of initiation factors IIA and IID and identification of a new factor operating at sequences downstream of the initiation site. J Biol Chem 262: 3322–3330. http://dx.doi.org/10.1016/S0021-9258(18)61506-6.

Robinson JT, Thorvaldsdóttir H, Winckler W, Guttman M, Lander ES, Getz G, Mesirov JP. 2011. Integrative genomics viewer. Nat Biotechnol 29: 24–26. http://www.nature.com/nbt/journal/v29/n1/abs/nbt.1754.html %5Cn http://www.nature.com/nbt/journal/v29/n1/pdf/nbt.1754.pdf.

Rossetti R, Ferrari I, Bonomi M, Persani L. 2017. Genetics of primary ovarian insufficiency. Clin Genet 91: 183–198.

Sangrithi MN, Royo H, Mahadevaiah SK, Ojarikre O, Bhaw L, Sesay A, Peters AHFM, Stadler M, Turner JMA. 2017. Non-Canonical and Sexually Dimorphic X Dosage Compensation States in the Mouse and Human Germline. Dev Cell.

Smale ST, Baltimore D. 1989. The “initiator” as a transcription control element. Cell 57: 103– 113.

Stuart T, Butler A, Hoffman P, Hafemeister C, Papalexi E, Mauck WM, Hao Y, Stoeckius M, Smibert P, Satija R. 2019. Comprehensive Integration of Single-Cell Data. Cell 177: 1888–1902.e21. https://doi.org/10.1016/j.cell.2019.05.031.

Szklarczyk D, Gable AL, Lyon D, Junge A, Wyder S, Huerta-Cepas J, Simonovic M, Doncheva NT, Morris JH, Bork P, et al. 2019. STRING v11: Protein-protein association networks with increased coverage, supporting functional discovery in genome-wide experimental datasets. Nucleic Acids Res 47: D607–D613.

Trapnell C, Cacchiarelli D, Grimsby J, Pokharel P, Li S, Morse M, Lennon NJ, Livak KJ, Mikkelsen TS, Rinn JL. 2014. The dynamics and regulators of cell fate decisions are revealed by pseudotemporal ordering of single cells. Nat Biotechnol 32: 381–386.

Verrijzer CP, Chen JL, Yokomori K, Tjian R. 1995. Binding of TAFs to core elements directs promoter selectivity by RNA polymerase II. Cell 81: 1115–1125.

Vizcaíno C, Mansilla S, Portugal J. 2015. Sp1 transcription factor: A long-standing target in cancer chemotherapy. Pharmacol Ther 152: 111–124.

Wickham H. 2016. ggplot2: Elegant Graphics for Data Analysis. 2nd ed. Springer-Verlag ggplot.org.

Yu G, Wang LG, Han Y, He QY. 2012. ClusterProfiler: An R package for comparing biological themes among gene clusters. Omi A J Integr Biol 16: 284–287.

Zhao ZH, Ma JY, Meng TG, Wang ZB, Yue W, Zhou Q, Li S, Feng X, Hou Y, Schatten H, et al. 2020. Single-cell RNA sequencing reveals the landscape of early female germ cell development. FASEB J 34: 12634–12645.

Zhou H, Grubisic I, Zheng K, He Y, Wang PJ, Kaplan T, Tjian R. 2013a. Taf7l cooperates with Trf2 to regulate spermiogenesis. Proc Natl Acad Sci.

Zhou H, Kaplan T, Li Y, Grubisic I, Zhang Z, Wang PJ, Eisen MB, Tjian R. 2013b. Dual functions of TAF7L in adipocyte differentiation. Elife.

