## Supplemental_Fig_1 for "TAF4b transcription networks regulating early oocyte differentiation"

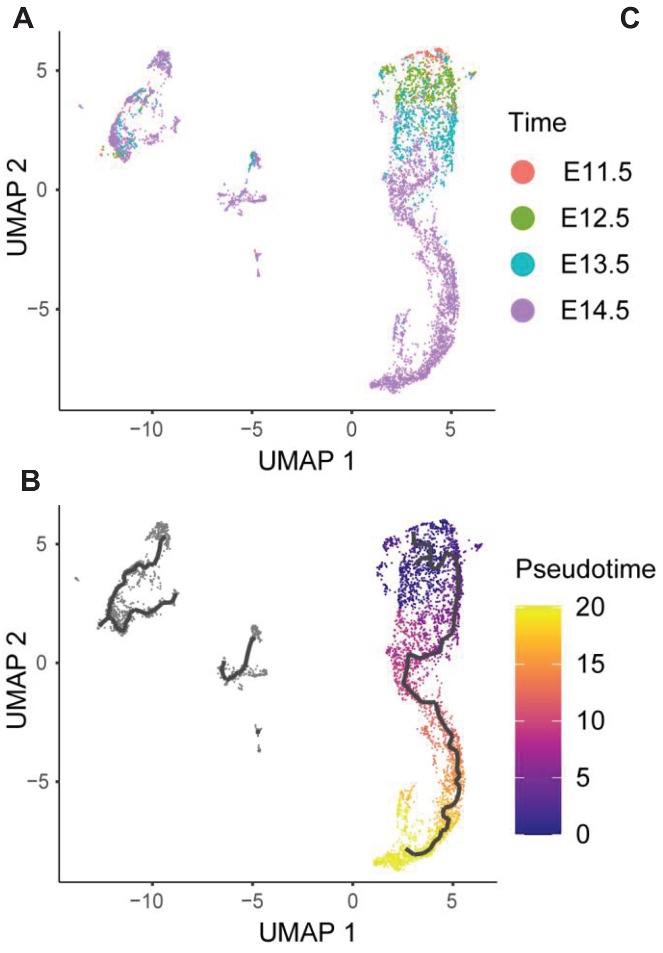


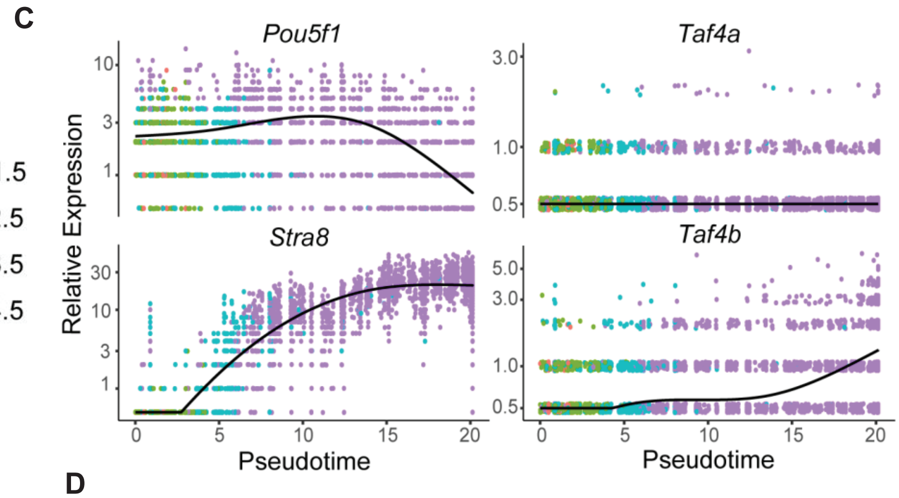


D

| Gene | Log2 Fold Change | P-Value |
|---|---|---|
| Taf4b | 11.78 | 4.34E-111 |
| Taf7l | 1.06 | 7.27E-04 |
| Tex12 | 0.98 | 1.71E-03 |
| Xlr4a | 0.98 | 2.13E-03 |
| Sycp3 | 0.97 | 2.40E-03 |
| Tex15 | 0.96 | 2.45E-03 |
| Tex101 | 0.90 | 5.62E-03 |
| Syce3 | 0.89 | 6.80E-03 |
| M1ap | 0.87 | 8.79E-03 |
| Meioc | 0.81 | 1.83E-02 |

GO Term

Synaptonemal Complex Assembly

Meiosis I/ Meiotic Cell Cycle

**Figure S1. Analysis of scRNA-seq dataset of E11.5 to E14.5 female germ cells.** (A) Uniform Manifold Approximation and Projection (UMAP) of oocytes colored by embryonic time point. (B) UMAP of oocytes colored by pseudotime analysis. (C) Expression of *Pou5f1*, *Stra8*, *Taf4b*, and *Taf4a* plotted in terms of pseudotime and colored based on embryonic time point. (D) Table of top 10 genes (in addition to *Taf4b*) that are expressed significantly higher in *Taf4b*-expressing oocytes. Colors indicate association with gene ontology (GO) term synaptonemal complex assembly (yellow) and meiosis I/meiotic cell cycle (red).
