## Supplemental_Fig_2 for "TAF4b transcription networks regulating early oocyte differentiation"

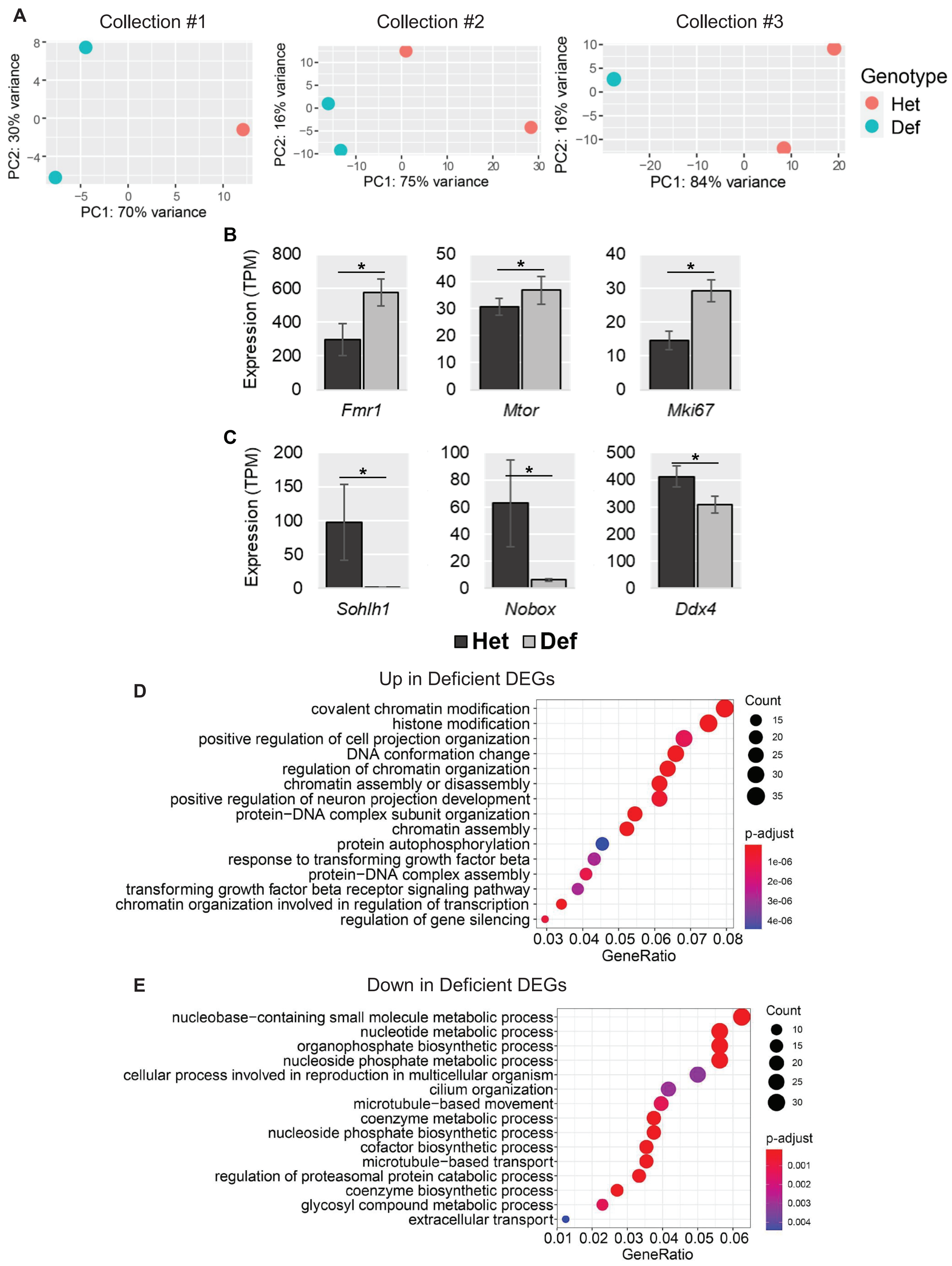

A
Collection #1
Collection #2
Collection #3
PC1: 70% variance
PC2: 30% variance
PC1: 75% variance
PC2: 16% variance
PC1: 84% variance
PC2: 16% variance
Genotype
Het
Def
B
Expression (TPM)
Fmr1
Mtor
Mki67
C
Expression (TPM)
Sohlh1
Nobox
Ddx4
Het
Def
D
Up in Deficient DEGs
covalent chromatin modification
histone modification
positive regulation of cell projection organization
DNA conformation change
regulation of chromatin organization
chromatin assembly or disassembly
positive regulation of neuron projection development
protein−DNA complex subunit organization
chromatin assembly
protein autophosphorylation
response to transforming growth factor beta
protein−DNA complex assembly
transforming growth factor beta receptor signaling pathway
chromatin organization involved in regulation of transcription
regulation of gene silencing
GeneRatio
Count
p-adjust
E
Down in Deficient DEGs
nucleobase−containing small molecule metabolic process
nucleotide metabolic process
organophosphate biosynthetic process
nucleoside phosphate metabolic process
cellular process involved in reproduction in multicellular organism
cilium organization
microtubule−based movement
coenzyme metabolic process
nucleoside phosphate biosynthetic process
cofactor biosynthetic process
microtubule−based transport
regulation of proteasomal protein catabolic process
coenzyme biosynthetic process
glycosyl compound metabolic process
extracellular transport
GeneRatio
Count
p-adjust

**Figure S2. E16.5 RNA-seq details.** (A) PCA plots of different E16.5 RNA-seq collections colored by genotype. (B) Expression levels of “up in deficient” DEGs in TPM. (C) Expression levels of “down in deficient” DEGs. (D) Biological process GO analysis dotplot of DEGs that were increased in *Taf4b*-deficient oocytes (“Up in Deficient”). (E) Biological process GO analysis dotplot of DEGs that were decreased in *Taf4b*-deficient oocytes (“Down in Deficient”).
