## Supplemental_Fig_3 for "TAF4b transcription networks regulating early oocyte differentiation"

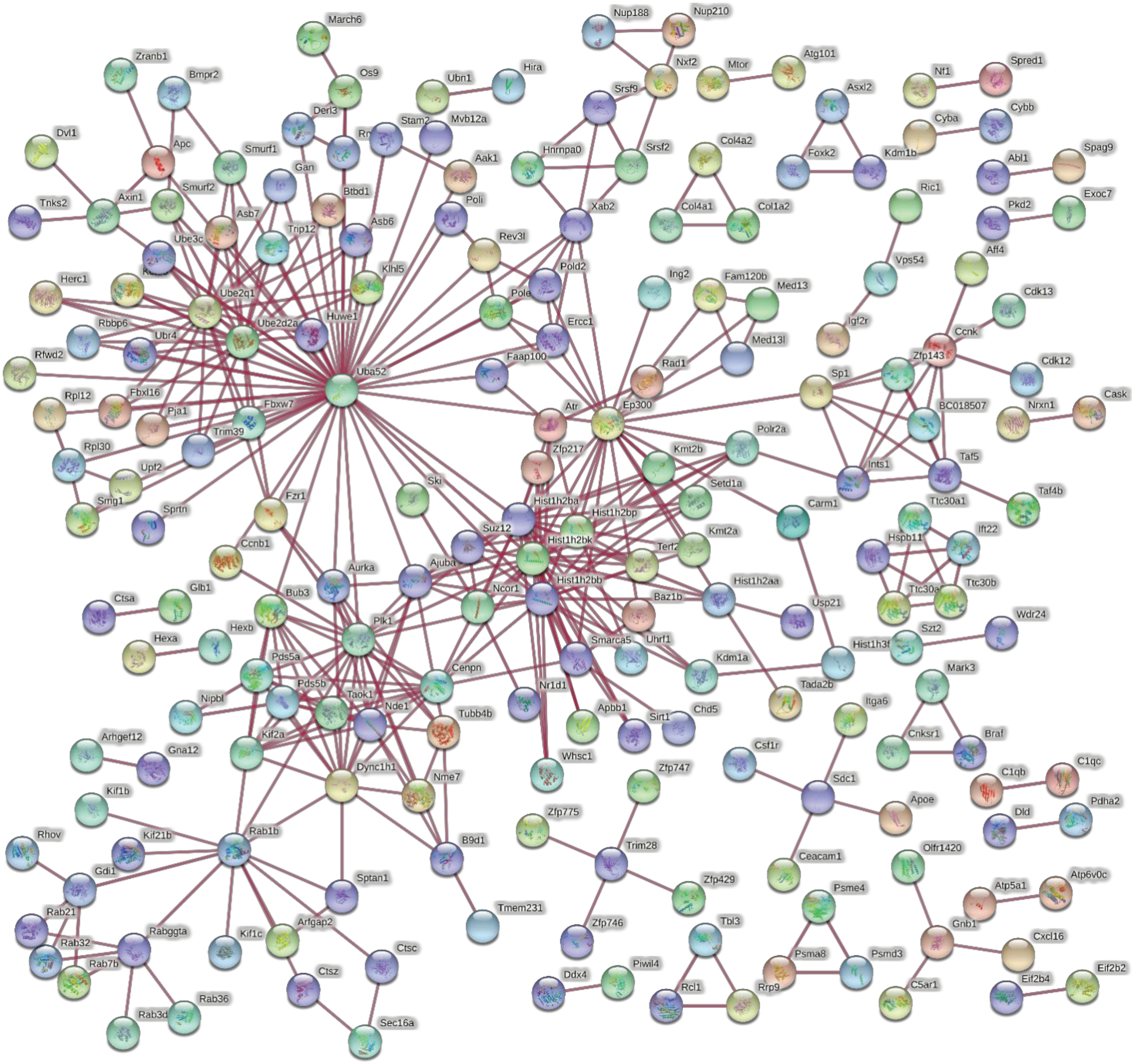


**Figure S3. Protein-protein interactions (PPIs) of E16.5 DEGs.** Plot generated by STRING of DEGs from E16.5 RNA-seq that had at least one PPI (physical network, highest confidence) (Szklarczyk et al. 2019). There was a significant enrichment of PPIs ($p < 0.01$).
