## Supplemental_Fig_4 for "TAF4b transcription networks regulating early oocyte differentiation"

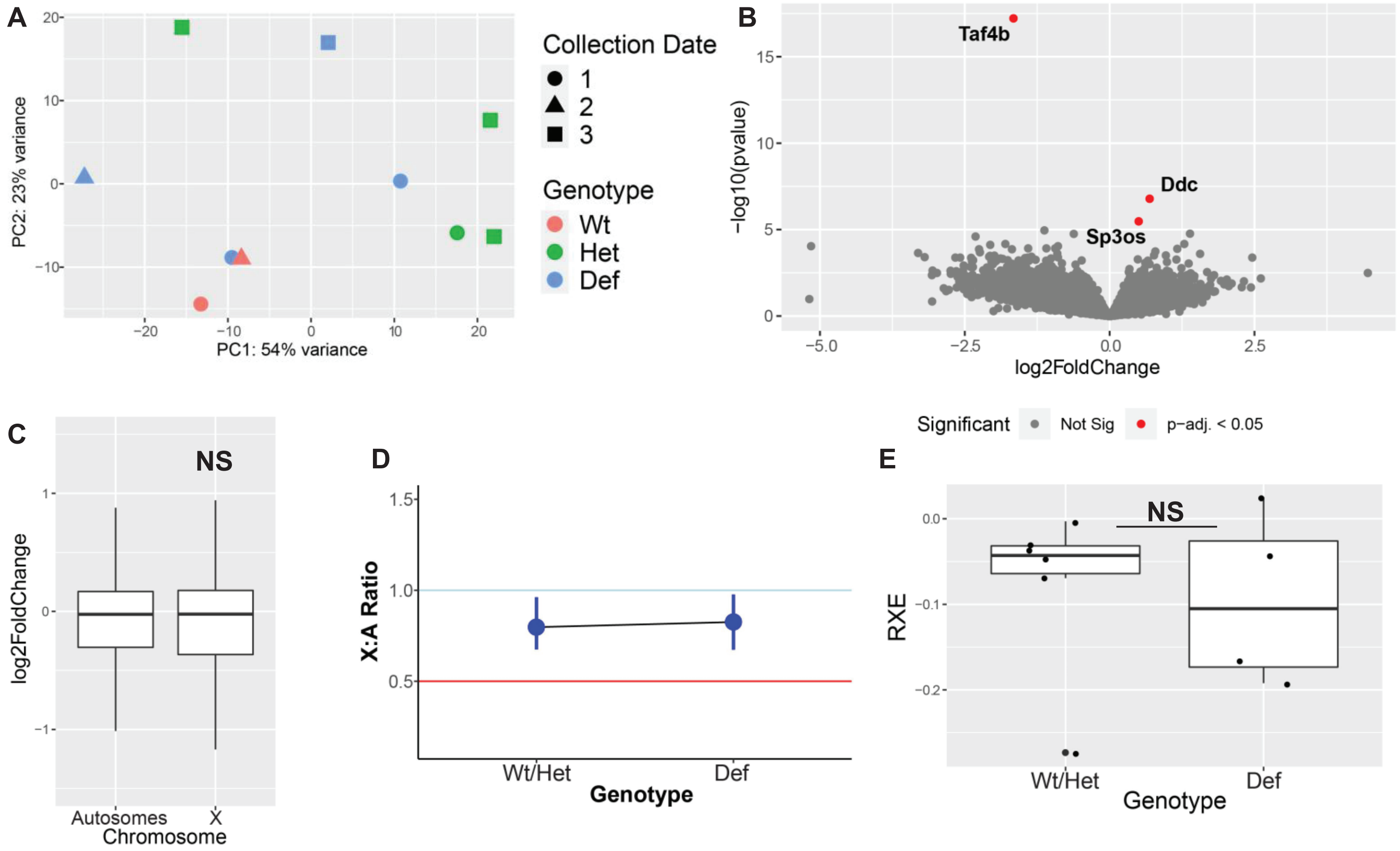


**Figure S4. E14.5 oocyte RNA-seq experiment.** (A) PCA of E14.5 RNA-seq with genotype and collection date plotted. (B) Volcano plot of E14.5 RNA-seq with DEGs labeled (red). (C) Boxplots of $\log_2$ fold change values from DESeq2 of all genes comparing autosomal versus X chromosome $\log_2$ fold change (outliers removed), NS = not significant. (D) X:A ratio plot calculated through pairwiseCI after filtering for average TPM > 1 comparing Wt/Het X:A ratio to *Taf4b*-deficient X:A Ratio. (E) Boxplots of relative X expression (RXE) calculations after filtering for average TPM > 1 and adding pseudocounts for log-transformation for each *Taf4b*-Wt, -heterozygous, and –deficient sample. Wt/Het samples compared to Defs, NS = not significant.
