## Supplemental_Fig5 for "TAF4b transcription networks regulating early oocyte differentiation"

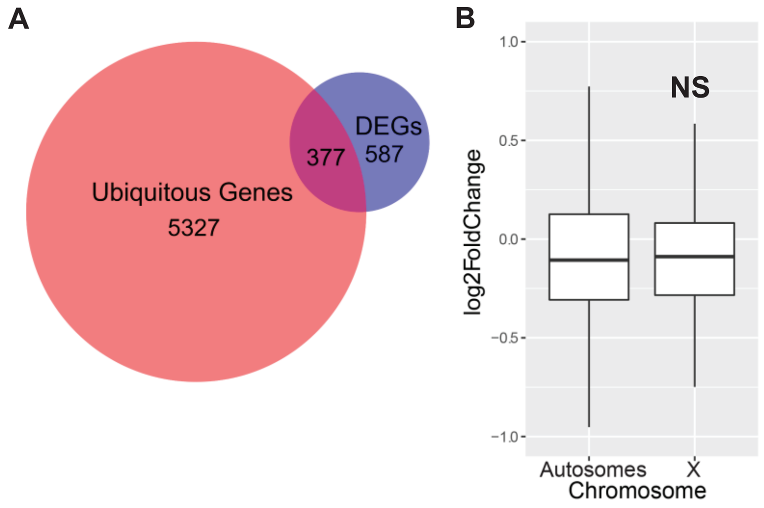


**Figure S5. Ubiquitous gene expression in E16.5 oocytes.** (A) Venn diagram of E16.5 DEGs and "Ubiquitous Genes" as identified in Sangrithi et al. 2017. Significant overlap in venn diagram ($p < 0.0001$, hypergeometric test). (B) Boxplots of $\log_2$ fold change values from DESeq2 of all ubiquitous genes comparing autosomal $\log_2$ fold change versus X chromosome $\log_2$ fold change, NS = not significant (outliers removed).
