## Supplemental_Fig_6 for "TAF4b transcription networks regulating early oocyte differentiation"

A

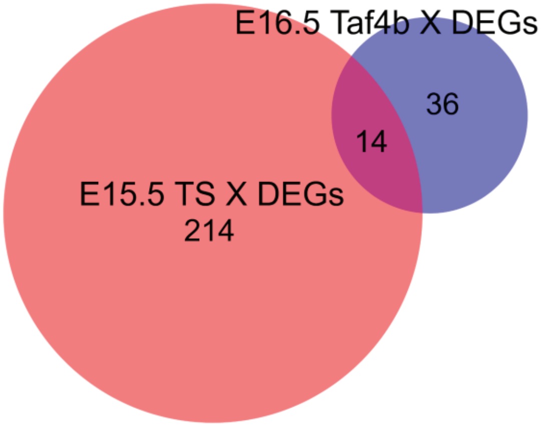


B

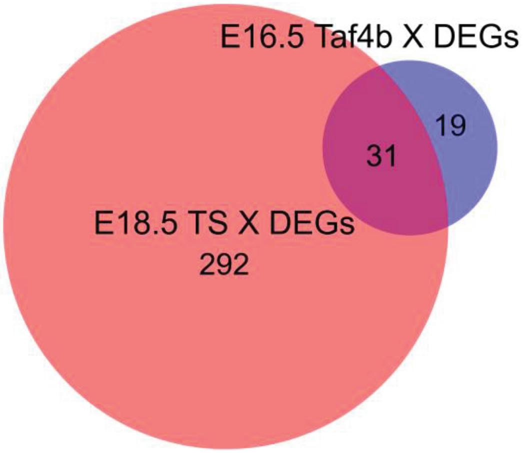


C

| Gene Name | Ensembl Gene ID |
|---|---|
| 4930550L24Rik (Magea13) | ENSMUSG00000046180 |
| Atg4a | ENSMUSG00000079418 |
| Ccdc160 | ENSMUSG00000073207 |
| Clcn5 | ENSMUSG00000004317 |
| Fmr1 | ENSMUSG00000000838 |
| Gdi1 | ENSMUSG00000015291 |
| Huwe1 | ENSMUSG00000025261 |
| Magea10 | ENSMUSG00000043453 |
| Mageb4 | ENSMUSG00000035427 |
| Nsdhl | ENSMUSG00000031349 |
| Nxf2 | ENSMUSG00000009941 |
| Rps4x | ENSMUSG00000031320 |
| Shroom4 | ENSMUSG00000068270 |
| Slc35a2 | ENSMUSG00000031156 |

D

| Gene Name | Ensembl Gene ID |
|---|---|
| 1700080O16Rik (Magea14) | ENSMUSG00000031118 |
| Atg4a | ENSMUSG00000079418 |
| AV320801 | ENSMUSG00000054994 |
| Bcorl1 | ENSMUSG00000036959 |
| Ccdc160 | ENSMUSG00000073207 |
| Clcn5 | ENSMUSG00000004317 |
| Cybb | ENSMUSG00000015340 |
| Fgf13 | ENSMUSG00000031137 |
| Fmr1 | ENSMUSG00000000838 |
| Gab3 | ENSMUSG00000032750 |
| Gdi1 | ENSMUSG00000015291 |
| Gla | ENSMUSG00000031266 |
| Gm15023 | ENSMUSG00000079432 |
| Gm364 (Tm9sf5) | ENSMUSG00000079584 |
| Gm5128 | ENSMUSG00000094004 |
| Gm7173 (Cfap47) | ENSMUSG00000073077 |
| Huwe1 | ENSMUSG00000025261 |
| Mageb18 | ENSMUSG00000035427 |
| Mageb4 | ENSMUSG00000035427 |
| Mbnl3 | ENSMUSG00000036109 |
| Nkrf | ENSMUSG00000044149 |
| Nsdhl | ENSMUSG00000031349 |
| Nup62cl | ENSMUSG00000072944 |
| Nxf2 | ENSMUSG00000009941 |
| Pja1 | ENSMUSG00000034403 |
| Rhox2d | ENSMUSG00000095698 |
| Rps4x | ENSMUSG00000031320 |
| Shroom4 | ENSMUSG00000068270 |
| Slc35a2 | ENSMUSG00000031156 |
| Tktl1 | ENSMUSG00000031397 |
| Tsga8 | ENSMUSG00000035522 |

**Figure S6. Shared X chromosome DEGs between TS dataset and *Taf4b*-deficiency.** (A) Venn diagram of all E16.5 *Taf4b* X chromosome DEGs compared with E15.5 TS X chromosome DEGs (protein-coding, p-adj < 0.05, avg TPM > 1). No significant overlap in Venn diagram (hypergeometric test). (B) Venn diagram of all E16.5 *Taf4b* X chromosome DEGs compared with E18.5 TS X chromosome DEGs (protein-coding, p-adj < 0.05, avg TPM > 1). Significant overlap in Venn diagram (p < 0.0001, hypergeometric test). (C) List of the 14 DEGs shared in (A). (D) List of the 31 DEGs shared in (B).
