## Supplemental_Fig7 for "TAF4b transcription networks regulating early oocyte differentiation"

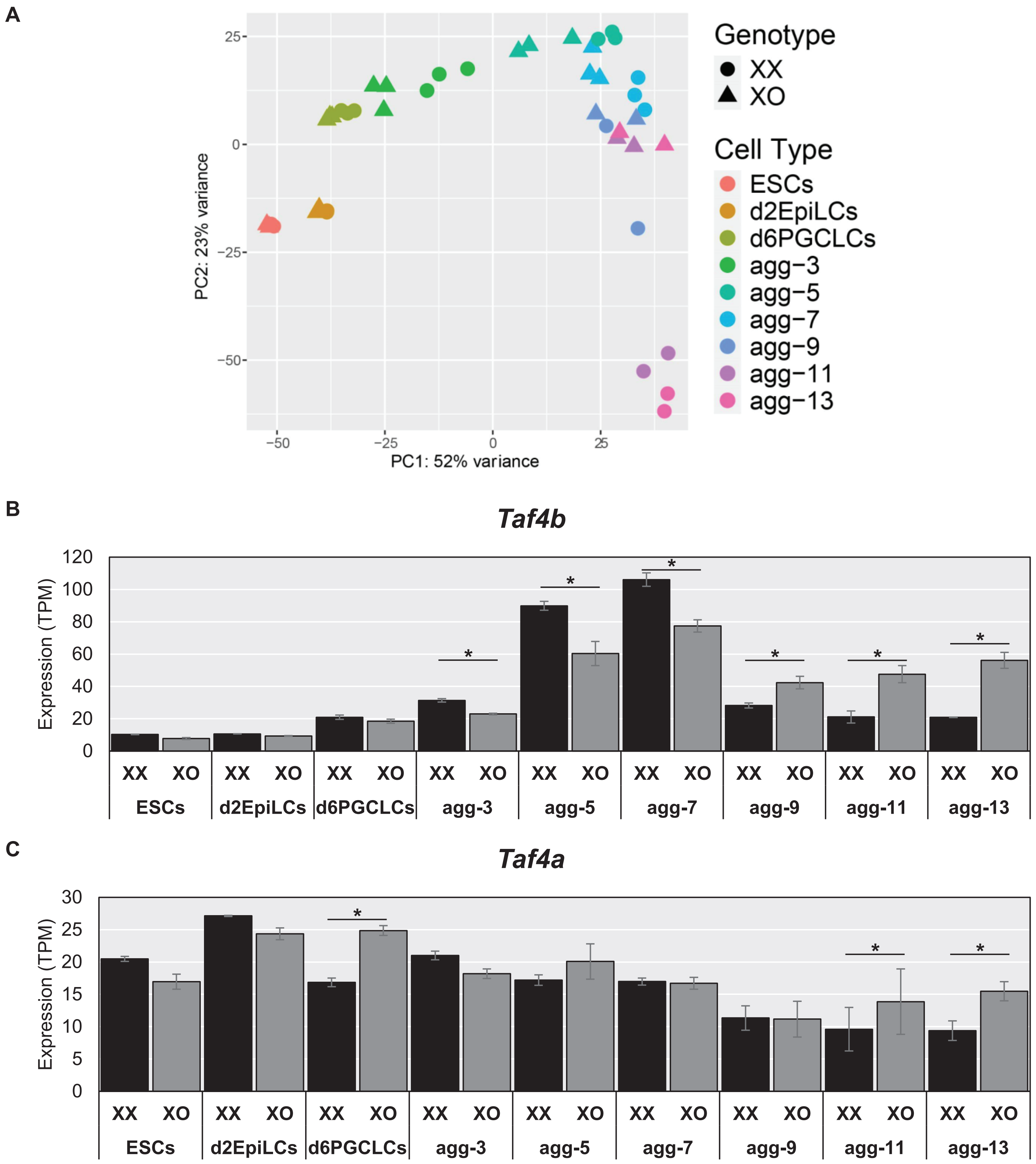


**Figure S7. Effects of XO on *Taf4b* in in vitro germ cell differentiation culture system.** (A) PCA plot of cultured cells from Hamada et al., 2020, labeled based on cell type and genotype. (B) Expression levels of *Taf4b* in XX versus XO cultured cells (* = p < 0.05, avg TPM > 1). (C) Expression levels of *Taf4a* in XX versus XO cultured cells. Error bars indicate ± standard error of the mean (SEM).
