## Supplemental_Fig_8 for "TAF4b transcription networks regulating early oocyte differentiation"

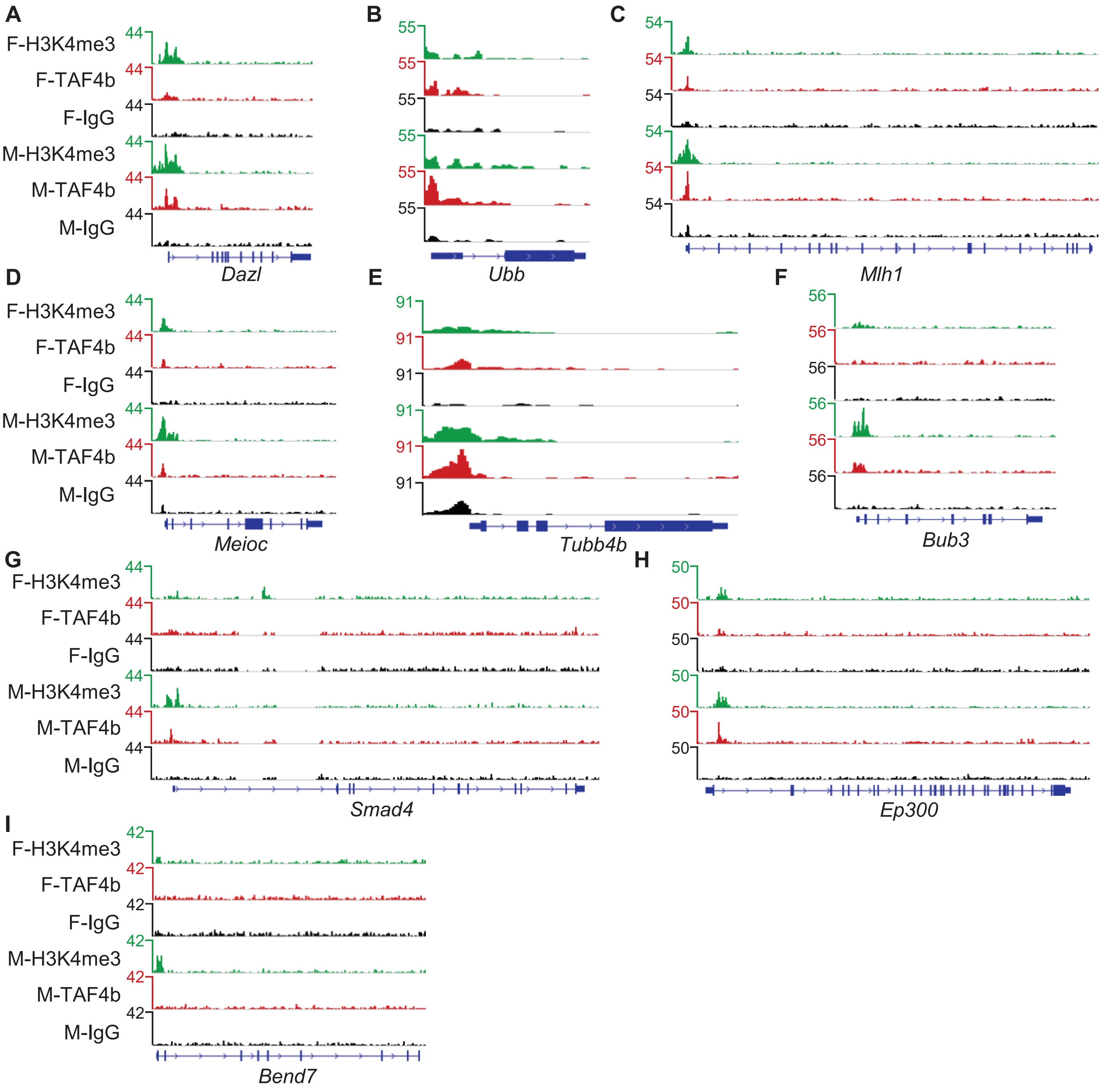


**Figure S8. Selected TAF4b-bound gene tracks.** (A) Gene track of *Dazl*, a shared CUT&RUN peak but not a DEG. (B) Gene track of *Ubb*, a shared CUT&RUN peak and a DEG. (C) Gene track of *Mlh1*, a female-only CUT&RUN peak and not a DEG. (D) Gene track of *Meioc*, a female-only CUT&RUN peak and not a DEG. (E) Gene track of *Tubb4b*, a female-only CUT&RUN peak and a DEG. (F) Gene track of *Bub3*, a male-only CUT&RUN peak and a DEG. (G) Gene track of *Smad4*, a male-only CUT&RUN peak and not a DEG. (H) Gene track of *Ep300*, a male-only CUT&RUN peak and a DEG. (I) Gene track of *Bend7*, a non-DEG that had no TAF4b peaks called but did have H3K4me3 peaks called for both female and male germ cells.
