## Supplemental_Table_4 for "TAF4b transcription networks regulating early oocyte differentiation"

**Table S4: Chromosome distribution of Down in Deficient DEGs**

| Chromosome | Total Genes | Observed | Expected | $Chi^2$ | p-value |
|---|---|---|---|---|---|
| 1 | 1439 | 32 | 32.41 | 0.006 | 0.9406 |
| 2 | 1630 | 35 | 36.71 | 0.086 | 0.7694 |
| 3 | 1178 | 21 | 26.53 | 1.217 | 0.2699 |
| 4 | 1290 | 28 | 29.05 | 0.04 | 0.8409 |
| **5** | **1499** | **20** | **33.76** | **6.014** | **0.0142** |
| 6 | 1145 | 17 | 25.79 | 3.158 | 0.0755 |
| 7 | 1750 | 33 | 39.42 | 1.135 | 0.2867 |
| 8 | 1116 | 25 | 25.14 | 0.001 | 0.9771 |
| 9 | 1281 | 34 | 28.85 | 0.975 | 0.3233 |
| 10 | 1039 | 27 | 23.40 | 0.581 | 0.4459 |
| 11 | 1592 | 38 | 35.86 | 0.138 | 0.7107 |
| 12 | 901 | 21 | 20.29 | 0.026 | 0.8722 |
| 13 | 940 | 26 | 21.17 | 1.156 | 0.2823 |
| 14 | 764 | 16 | 17.21 | 0.088 | 0.7666 |
| 15 | 825 | 23 | 18.58 | 1.092 | 0.296 |
| 16 | 655 | 11 | 14.75 | 0.982 | 0.3216 |
| 17 | 1056 | 21 | 23.78 | 0.341 | 0.5591 |
| 18 | 571 | 15 | 12.86 | 0.365 | 0.5455 |
| 19 | 673 | 17 | 15.16 | 0.23 | 0.6313 |
| **X** | **900** | **41** | **20.27** | **22.094** | **<0.0001** |
