## Supplemental_Table_5 for "TAF4b transcription networks regulating early oocyte differentiation"

**Table S5: Chromosome distribution of Up in Deficient DEGs**

| | | | Expected | Chi$^2$ | p-value |
|---|---|---|---|---|---|
| 1 | 1439 | 24 | 29.95 | 1.264 | 0.2609 |
| 2 | 1630 | 40 | 33.93 | 1.172 | 0.279 |
| **3** | **1178** | **14** | **24.52** | **4.766** | **0.029** |
| 4 | 1290 | 30 | 26.85 | 0.392 | 0.5311 |
| 5 | 1499 | 38 | 31.20 | 1.589 | 0.2074 |
| 6 | 1145 | 18 | 23.83 | 1.504 | 0.2201 |
| 7 | 1750 | 40 | 36.43 | 0.381 | 0.5372 |
| 8 | 1116 | 21 | 23.23 | 0.225 | 0.6351 |
| 9 | 1281 | 28 | 26.66 | 0.071 | 0.7898 |
| 10 | 1039 | 18 | 21.63 | 0.638 | 0.4245 |
| **11** | **1592** | **48** | **33.14** | **7.181** | **0.0074** |
| 12 | 901 | 24 | 18.75 | 1.529 | 0.2162 |
| **13** | **940** | **31** | **19.57** | **6.977** | **0.0083** |
| 14 | 764 | 10 | 15.90 | 2.269 | 0.132 |
| 15 | 825 | 14 | 17.17 | 0.609 | 0.4353 |
| 16 | 655 | 9 | 13.63 | 1.623 | 0.2027 |
| 17 | 1056 | 19 | 21.98 | 0.424 | 0.5148 |
| 18 | 571 | 12 | 11.89 | 0.001 | 0.9731 |
| 19 | 673 | 16 | 14.01 | 0.292 | 0.5889 |
| **X** | **900** | **9** | **18.73** | **5.268** | **0.0217** |
