## Supplemental_Table_10 for "TAF4b transcription networks regulating early oocyte differentiation"

**Table S10: Cell numbers for RNA-seq samples**

| Age | Genotype | Sample # | Cell # |
|---|---|---|---|
| E14.5 | Wildtype | 1 | 7,942 |
| | | 2 | 18,256 |
| | Heterozygous | 1 | 12,553 |
| | | 2 | 12,897 |
| | | 3 | 2,822 |
| | | 4 | 19,308 |
| | Deficient | 1 | 9,112 |
| | | 2 | 5,233 |
| | | 3 | 9,893 |
| | | 4 | 7,035 |
| E16.5 | Heterozygous | 1 | 7,199 |
| | | 2 | 2,512 |
| | | 3 | 3,399 |
| | | 4 | 9,369 |
| | | 5 | 14,402 |
| | Deficient | 1 | 2,181 |
| | | 2 | 3,547 |
| | | 3 | 19,089 |
| | | 4 | 5,688 |
| | | 5 | 9,076 |
